# Learning causal regulatory motifs and grammars using deep learning models and massively parallel reporter assays

**DOI:** 10.1101/2025.07.25.666754

**Authors:** Mike Thompson, Ben Lehner

## Abstract

A central challenge in biology is to understand, predict, and engineer the ‘second genetic code’: how sequence encodes gene expression. Two components of this challenge are: (1) accurate prediction (and design) of gene expression from sequence and (2) mechanistic understanding of how sequence-to-expression encoding actually works in cells. A powerful general approach to this problem is to combine large scale data generation with artificial intelligence. For example, massively parallel reporter assays (MPRAs) can quantify the expression of thousands of different sequences in pooled experiments and the resulting data can be used to train deep learning models. Unlike in the case of long-context genomic language models, where transformer-based architectures are a dominant paradigm, it remains contested whether for MPRA datasets other architectural components can lead to more useful, generalizable predictors, and whether they affect model interpretability, i.e. the ability to capture causal biological mechanisms (either inherently or when using downstream interpretability or explainability techniques, “xAI”). Ablation analyses may help elucidate important architectural components, but are almost always anecdotal, unable to describe generalizable tendencies, as they are done with a single training dataset or a few testing datasets. Here, we attempt to reconcile concerns and provide guidance for MPRA model design and xAI choice by simulating at scale 1,500 motif-based genetic architectures and evaluating the ability of different model architecture-xAI pairs to first predict an outcome given a sequence as input, and second, report involved motifs and their corresponding grammar. We find that attention-based models are efficient learners, and while we recommend their use in low-data regimes, their performance is surpassed by alternative models, like dilated CNNs, under larger sample sizes. We next show that across grammars and models, current methods for motif extraction converge toward reporting the same set of motifs, which is dominated by motifs with large effect sizes. We then perform *in silico* experiments across models and their discovered motifs and find that these methods accurately rank motifs based on learned effect size, but that their learned effect size is systematically miscalibrated, particularly in the presence of interactions (epistasis). Finally, we propose a novel metric for identifying motifs involved in epistasis and confirm our findings across three experimental datasets. Our work provides practical guidance for modeling and interpreting massively parallel reporter assay experiments from end to end.

## 1 Introduction

Massively parallel reporter assay (MPRAs)—in which a biological sequence is systematically altered and its downstream function measured—hold the potential for not only clarifying variant effects on disease, but also for elucidating novel functional or mechanistic insight[1, 2]. In recent years, these experiments have moved beyond simple characterization of genomic sequences and single-nucleotide mutational effects to also include more complicated modifications—higher order mutations[3], insertion/deletions[4], shuffling[5, 6], and randomization of sequences[7,8,9,10]. Nonetheless, as these experiments have increased in both scale and complexity, the need for model-based assistance to glean insight from these datasets has too grown.

Neural networks have emerged as the tool of choice for modeling MPRA experiments owing to their state-of-the-art predictive performance across a wide variety of regulatory processes[7, 9, 11, 12, 13, 14, 15]. Despite there remaining debate about whether these models are interpretable, either inherently, or when using downstream explainability techniques (xAI), there has been substantial progress made in extracting model-learned representations and grammar[15, 16]. Currently, interpretability analyses typically involve (1) generating some form of position-variant score, (2) identifying repetitions of highly important sub-strings of a sequence (e.g., motifs), and (3), performing *in silico* analyses where the model is treated as a proxy for the experiment, and mechanistic hypotheses are probed by modifying sequences and evaluating the model’s predicted changes on function[14,17, 18]. While this pipeline has become the *de facto* field standard, there remains much uncertainty of whether certain genetic grammars, neural network architectures, importance scores, or combinations thereof, are more amenable to discovering or faithful to revealing the underlying biology than others.[14,15,16].

In terms of architectures, neural networks for MPRA data have leveraged convolutions as the near-universal first-layer architectural component since their introduction into to the field[11, 12, 13, 14, 19]. This is primarily due to two reasons: first, allowing a model to learn its own features for prediction tasks is a rational design choice, and second, convolutions have an immediately reifiable meaning as position weight matrices (PWM), the dominant data structure of biological motifs. Optimal selection of downstream components of architectures, on the other hand, remains less trivial. In recent years, attention (or transformer) blocks have gained substantial traction as the dominant architectural paradigm, given their computational efficiency, wide field-of-view, as well as their ubiquity and successes in large language model (LLM) approaches[20, 21]. Despite this, several state-of-the-art task-specific models of DNA regulatory processes—e.g. of cell type-specific acessibility [22] or splicing—[23] leverage stacked convolutions, or dilated convolutions, and in several cases outperform more complex, multi-headed, U-net or deep attention-driven approaches (e.g. in comparisons with Enformer[22, 24], or certain splicing tasks with SpliceAI and AlphaGenome[25]). Why these architectural-based differences in performance occur and whether such differences would be meaningful in the context of modeling short sequences—i.e., whether they are indicative of learning specific biological grammars, artifacts of sample sizes, or other sources of variability—remains understudied.

While powerful attribution and motif discovery tools exist[26,27,28], the field lacks systematic guidance on which combinations of architectures and xAI methods yield reliable motif discovery under which genomic grammars[14,17, 18]. There have been promising early explorations of xAI methods and architectures on a small set of simulated dat 29], however most existing evaluations consider few (*<* 10) realizations of specific biological or simulated systems and neural network architectures [14, 18, 30,31]. We believe these types of architectural or ablation-style comparisons are inherently anecdotal, in that they tend to capture differences in performance of architectures on only a *single* realization of a specific genomic grammar, i.e., experimental dataset. Indeed, evaluating out-of-sample performance on test-data can help fortify observations, but there is a critical dearth of comparisons of generalizable, long-term (i.e., statistical) tendencies of model architectures across many realizations of specific genetic grammars—something which is not ameliorated by evaluating on a training dataset and several out-of-sample test datasets. To clarify, we can imagine a hypothetical scenario in which MPRA data is collected for 500,000 sequences in a genomic context where functional readout is specified by some noise and a coordination of mostly additive-effect motifs with one simple epistatic interaction whose effect is much weaker than the additive effects. Training a variety of models on this data—including CNNs, transformers, long short-term memory (LSTMs)—and performing ablations on the corresponding optimal choice—pooling, dilations, etc.—can only reveal that the architecture choice was optimal under this specific realization of this genomic grammar, noise, and sample size. In other words, this single dataset is not sufficient to suggest that one architecture or another, is, in general, more adept at broadly representing genomic grammars (including those that are purely epistatic, purely additive, or both), nor under what sample sizes. Returning to the example, it could be that the attention module excels at this specific realization of data, but is surpassed by dilated CNNs under the same grammar, but when the epistatic effect is only slightly smaller than the additive effects. Without knowing the general tendencies of model architectures, researchers wishing to model their own datasets are faced with the seemingly un-answerable question of how they should model their data, and whether the differences that they observe in model performance are expected, or even meaningful.

In this work, we investigate how genetic grammar, network architecture, importance score methods, and design of *in silico* experiments influence prediction performance and interpretability insights by performing 1,500 simulation experiments and evaluating our findings on real, external MPRA datasets. By performing our simulations at scale, we are able to propose general tendencies, rather than realization-specific anecdotes, which we validate in orthogonal, biological experiments spanning both random and genomic contexts. We are hopeful that our experiments here guide and ameliorate concerns over architecture and xAI choice when modeling short-context (~200-300 basepair) MPRA data, and we summarize our findings and guidelines as follows:

- **Attention is an efficient learner**. Attention excels at prediction tasks in the short-sequence context under low sample sizes, but is surpassed by more complicated architectures (dilated CNNs, LSTMs) given sufficient data. We accordingly recommend attention in settings with low sample sizes.
- **Importance scores vary in motif preferences but converge toward the same set of motifs**. Across attribution methods, including gradients, *in silico* mutagenesis (ISM) and more-involved approaches, the set of discovered motifs is quite stable, and is dominated by motifs with large effect size. We suggest using (corrected[32]) gradients as a trade-off of speed and comprehensiveness for attention-based models and ISM for all other architectures due to its robustness. Visualizing first-layer filters can offer additional information when trained over random background sequences.
- ***in silico* approaches for interpreting motif effect size rank well but are miscalibrated**. xAI methods exhibit strong correlation between true and learned effects, but are miscalibrated depending on the network architecture and whether epistasis or sequence-based, non-motif effects are present. We therefore posit that probed effect sizes can be trusted to rank motif importance.
- **Second-moment information can detect motifs with epistatic effects, but their detection remains labor-intensive**. Here, we suggest a novel metric for predicting whether a motif is involved in a model-learned interaction (which is linear in the number of motifs, and therefore more efficient than current methods which are generally quadratic in the number of motifs), but show that power is modest. We recommend caution when interpreting epistasis, as signal may be spuriously due to simple main-effect miscalibration or non-motif background effects.

## 2 Results

We carried out from start to finish the current state-of-the-art MPRA deep-learning workflow on over 1,500 dataset instances, where, for each instance, we trained 8 neural network architectures, generated multiple attribution scores per architecture, extracted learned motifs per model-attribution pair, and finally, ran *in silico* experiments to probe per-model learned grammar of discovered motifs. We began our analyses with simulations based on a variance components model that enabled us to vary for each instance the additivity of motifs, epistasis of motifs, magnitude of additive and epistatic effects, sample size, and finally, interaction mechanism of each motif (Methods). After drawing conclusions from our comprehensive simulated datasets, we validated our findings on real MPRA experiments comprising first, a dataset measuring functional activity of random, synthetic sequences, and second, two that measured functional activity in endogenous, genomic-context sequences across cell-types.

### Attention is an efficient learner

We begin our analyses with a series of simulated MPRA experiments to elucidate expectations and tendencies of model architecture predictive performance in short sequence-to-phenotype prediction tasks across a variety of parameters and contexts. We varied the simulations to include multiple dataset sizes (few or many sequences) and genetic architectures, ranging from scenarios in which motif effects are exclusively additive to exclusively interactive, as well as scenarios in which there exist non-motif sequence effects that explain varying levels of phenotypic variance (Methods). Importantly, all models shared the same first layer—comprising a convolution with exponential activation [37]—as it has been widely shown to serve as a powerful inductive bias for both strong prediction accuracy and interpretability. For deeper layers, we employed a collection of architectures that are prevalent in the genomics space, including stacked convolutions[13], stacked dilated convolutions[23, 35], attention layers[33, 34], and long short-term memory (LSTM) units[36, 38], all prior to passing their constructed representations into a shallow multi-layer perceptron (MLP; Table 1). We also evaluated whether certain components of these architectures (e.g. skip/residual connections[39]) had significant effects on the predictions made by the models.

**Table 1:**
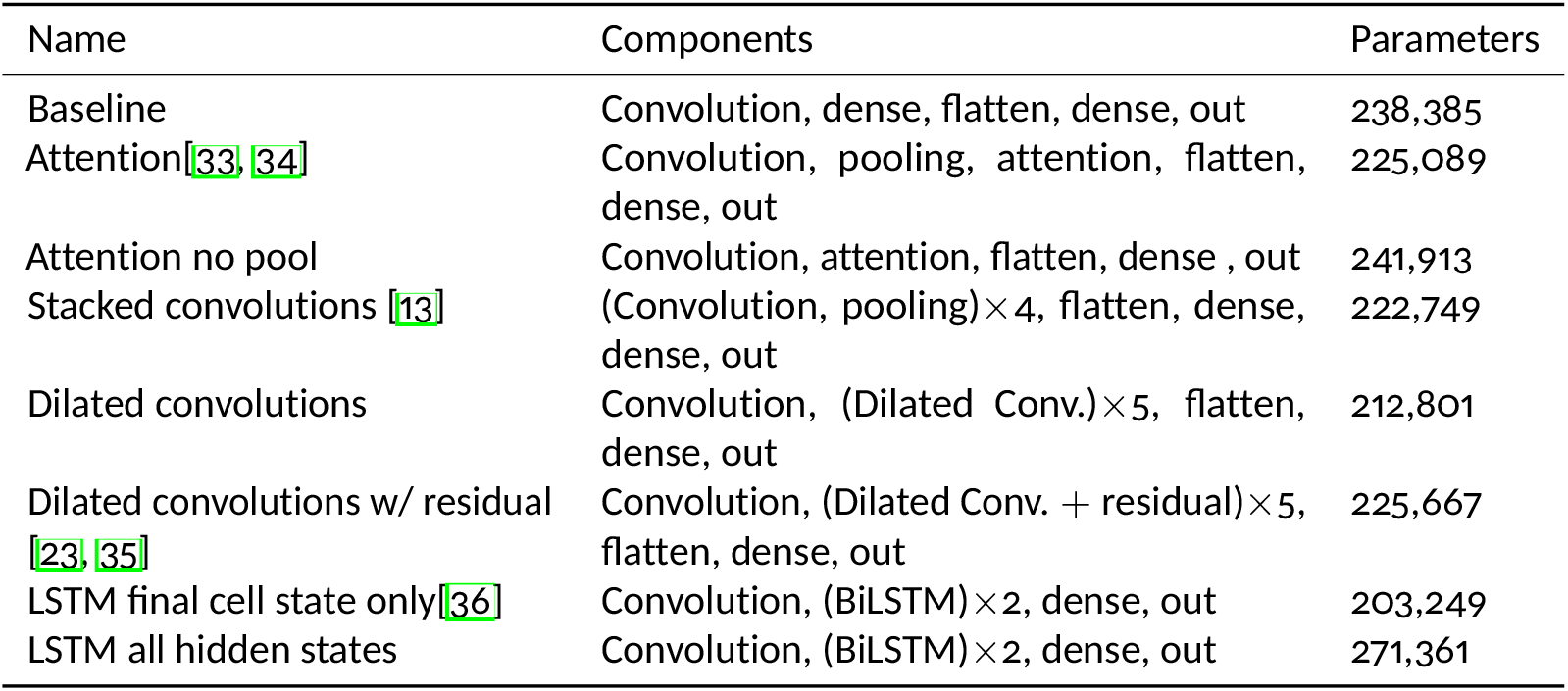
Architectures evaluated in the simulations. In this work, we set as a baseline inductive bias a motif detector (first-layer convolution with exponential activation), and compare, under a similar parameter budget, the efficacy of deeper-layer architectural components to produce useful features before a final MLP output head. Additional details are in the Methods section.

For each genetic architecture-sample size pair, we generated 50 unique datasets (each with their own realizations of motif frequency, effect size and epistasis), fit a model on 90% of the dataset, and evaluated model performance on the remaining 10% (Fig. 2, Supp. Figs.S1,S2). Across all tested genomic grammars, models that used attention blocks substantially outperformed other architectures in the low-sample-size regime, with differences becoming more pronounced as difficulty of genomic grammar increased (percent improvement over next best model in median predictive accuracy = 21.7% in the entirely additive context, 41.5% in the half-additive half-epistatic context, and 101.7% in the entirely-epistatic context). Once given 100,000 sequences, this disparity shrunk, with other architectures even out-performing attention-based models when non-additive effects were present (improvement in median score under pure motif additivity = 0.9%, decrease in accuracy of −3.7% in the half-additive half-epistatic context, and −8.3% in the fully epistatic context). When given 300,000 sequences, models using attention components were dominated by other model architectures, with the highest-performing model (using LSTMs) surpassing attention by 7.5% in the additive setting, 9.3% in the half-additive half-epistatic setting, and 13.6% in the entirely epistatic setting. We emphasize that, while architectures using convolutions and LSTMs were as competitive or more powerful than models using attention in high sample-size settings, the differences between these models and attention-based models were more-or-less negligible when the underlying genetic grammar was entirely additive.

**Figure 1:**
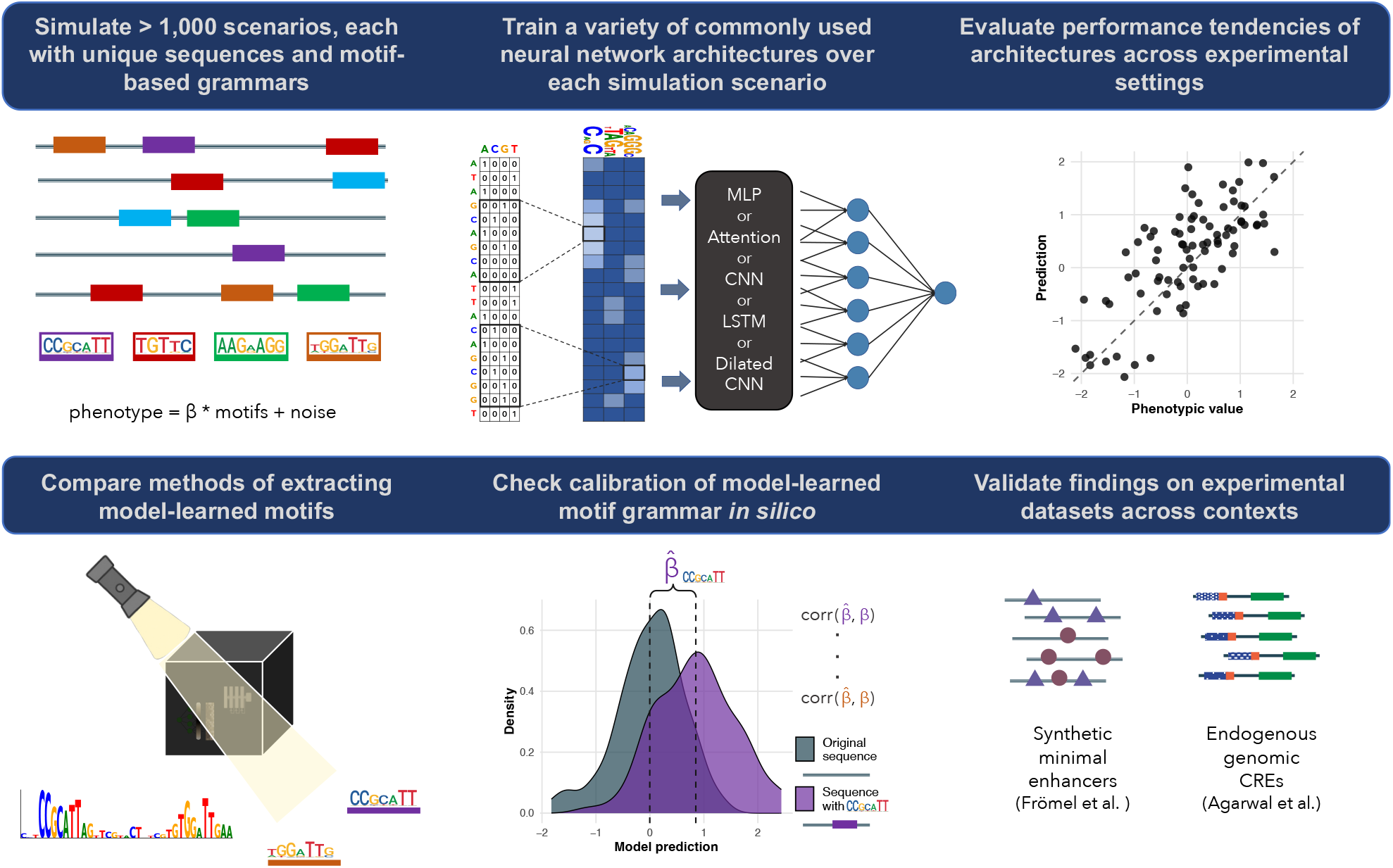
A systematic evaluation of neural network architectures and explainable AI methods across experimental and genomic contexts. We carry out from start to finish the current state-of-the-art deep learning pipeline for MPRA data on 1,500 simulated datasets under a motif-based generative model. We simulate sequences, motifs, and phenotypes under the assumption that the measured phenotype is explained by additive and/or epistatic motif effects and some noise (e.g., experimental noise). Next, we fit a variety of commonly used neural network architectures, compare their predictive accuracies, and perform a series of explainable AI (xAI) analyses to make sense of model-learned grammar. Finally, we validate our findings on three real, experimental datasets.

**Figure 2:**
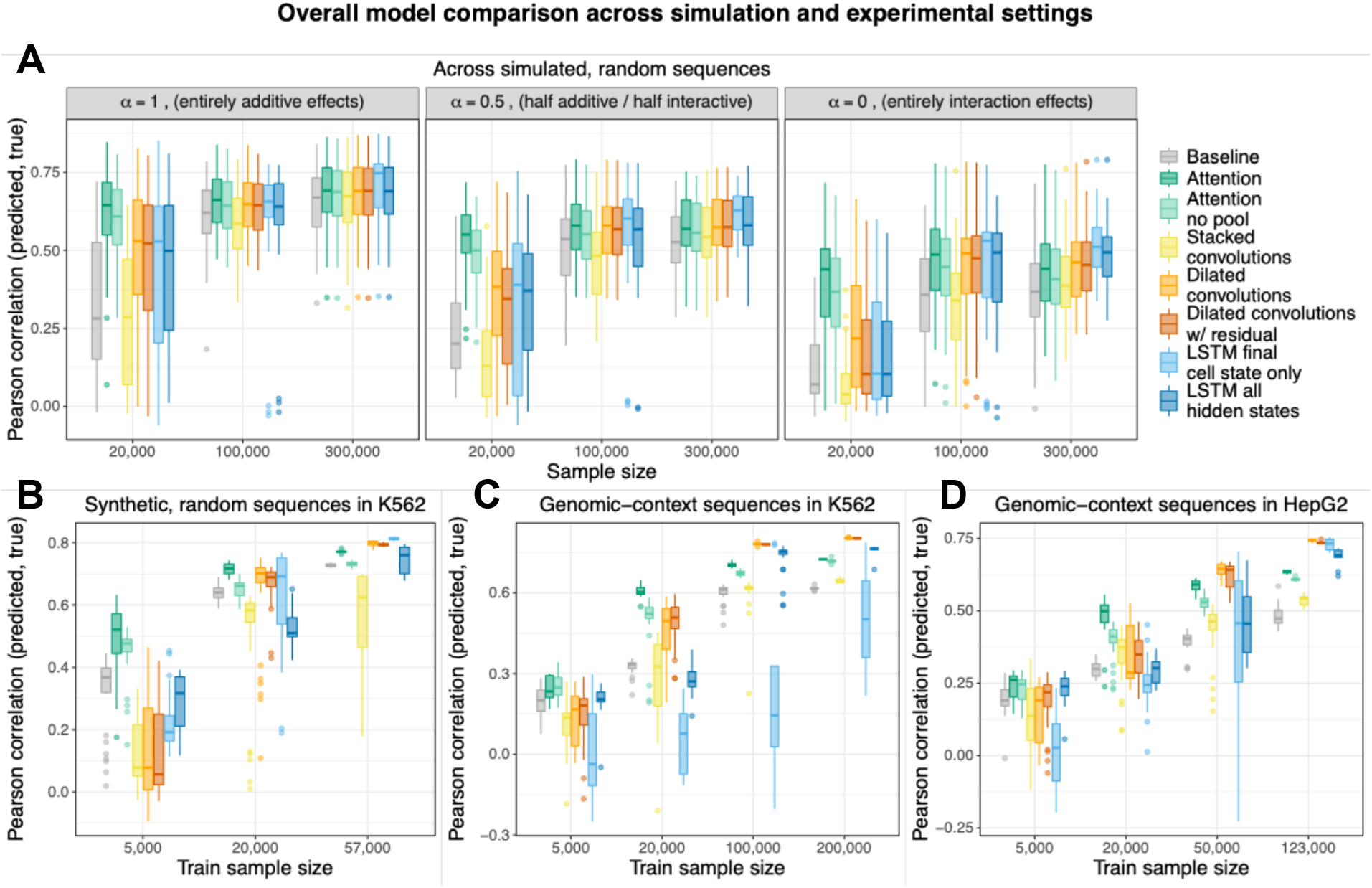
Genetic architectures and data availability modulate network performance. (A) Pearson correlation between the predicted and true phenotype across the 10% heldout simulated data. Each point underlying a boxplot represents a correlation calculated from a single training instance of a model under one specific genetic architecture and sample size configuration. For the simulated data, we fit one model across 50 different realized genetic architectures (e.g. varying motif presence, effect size, interaction effects, etc. See Methods; each boxplot comprises 50 points, and therefore genetic architecture realizations under a given sample size). (B-D) The performance of each model when training and predicting on a heldout proportion data for (B) the random, synthesized sequences of Frömel et al. [40] in K562 cell-lines, (C) the genomic-context sequences of Agarwal et al. [41] in K562 cell-lines, and (D) the genomic-context sequences of Agarwal et al. [41] in HepG2 cell-lines. In (B-D), the right-most x-axis entry represents the total dataset size, where all other entries were downsampled from this full dataset. Boxplots represent the distribution of 10-training instances for a given model-downsampled-data pair, where for a given downsampled dataset, the data were randomly sampled from the full data 3 times (for a total of 3 × 10 = 30 points in each boxplot).

To better understand the components underlying differences in model prediction accuracy, we also trained ablated versions of models—for the attention model, we tested the effect of pooling after the first convolutional layer; for the CNNs, we tested the effect of using dilations or not, as well as the utility of residual/skip connections in the dilated blocks; for the LSTMs we tested whether the hidden states at each position or the final cell-state were more informative. Attention models that used pooling significantly outperformed those that did not (paired t-test of out-of-sample Pearson correlations across sample-sizes and motif-additivity p*<*2.2e-16, mean difference=0.02). Though the differences were not significant at lower sample sizes, LSTMs that used only the final cell-state were significantly more accurate than those that used the hidden states along the sequence under large sample sizes, suggesting that given enough data, the architecture can learn to compress its modeled phenotype into a more compact vector (paired t-test over 150 Pearson correlations at N=300,000 p=0.003, mean of differences = 0.008). Under lower sample sizes, CNNs benefitted substantially from dilations, suggesting that having access to the entire sequence (or composite features thereof) is critical to accurate modeling, likely because the network cannot properly construct composite, hierarchical features from so few examples (t-test between CNN correlations and the pooled correlations from both dilated CNNs at N=20,000: t=-7.97 p=2.13e-14 and N=300,000: t=-2.33 p=0.02). This is further evidenced by the fact that under large sample sizes (N=300,000) and when the genetic grammar was entirely additive (and therefore no need to combine motifs), there were no significant differences between CNNs with and without dilations (t=-0.76, p=0.45), however in the fully epistatic grammar there were (t=-2.43, p=0.02). Finally, we note that by forcing the model to include the sequence in its learned representations, residual/skip connections in the dilations can potentially add noise to the modeling process when there are non-additive effects in the grammar (which depend on composite, higher-order features), as evidence by the discrepancies in dilated CNN performance in the presence of epistatic effects at low sample sizes that are ameliorated under high sample sizes (N=20,000 and entirely additive paired t-test of correlations between dilated CNNs with and without residual/skip connections: mean of differences=-0.01 p=0.77, and entirely epistatic: mean of differences=-0.06 p=4.1e-04; and N=300,000 entirely additive: mean of differences=-0.001 p=0.005; entirely epistatic: mean of differences=0.004 p=0.07)

We verified our findings by training the same 8 models on a collection of three MPRA datasets we briefly describe below. The first, of Frömel et al, comprised synthesized random sequences with varying combinations of transcription factor motifs and their measured downstream transcriptional activity in K562 cells (N=57,342, length=256; Methods)[40]. The second and third datasets comprised endogenous, human genomic cis-regulatory elements (CREs) assayed in K562 and HepG2 cell-lines (N=196,664 and 122,926 respectively, length=230; Methods) [41]. Unlike the sequences of Frömel et al, which consist of synthetic random sequences with implanted motifs, these datasets contain native genomic sequences, and therefore preserve genomic sequence context.

On these datasets, we performed a downsampling experiment whereby we downsampled from the original dataset under 3 different seeds and trained 10 model instances per each architecture and downsample split (i.e., for a given downsampled dataset, we trained 3 × 10 = 30 models for each network architecture). Our results were nearly entirely consistent with our simulations (Fig. 2). Primarily, models that used attention were most powerful in the low-sample-size setting (increase in median prediction accuracy ranged from 9.4-41.6%), but were later surpassed by dilated CNNs (dilated CNN increase in median prediction accuracy over attention-based models ranged from 3.8-17.0%). Interestingly, in the dataset of Frömel et al., which comprised random, synthetic sequences with injected motifs (most alike our simulated data), the LSTM architectures surpassed all other architectures, whereas in the genomic-context sequences of Agarwal et al., LSTMs generally failed to converge and their performance never seemed to plateau or surpass other architectures. Moreover, compared to models trained on random-context sequences, models trained on genomic-context sequences generally required many more samples to reach strong predictive accuracy, implying the genomic-context grammar is either more complicated, or simply harder to model, as sequence artifacts add noise to the underlying motif-based grammar.

### xAI methods and importance scores converge toward the same set of motifs

We next evaluated models’ abilities to capture the causal motif for all trained models and genetic architectures. We note that there exists a multitude of ways in which to extract motifs from a trained model. Here, we focus on two of the most prevalent methods—visualizing first-layer convolutional filters and running TF-MoDISco [28] over model attribution maps for a variety of importance scores, including gradients (×input), *in silico* mutagenesis (ISM)[13], and Shapley values (“SHAP”; Methods)[26]. Intuitively, the first-layer filters approach should reveal features that are recognized and either directly used to predict the phenotype, or that are non-trivially combined in deeper layers to predict the phenotype[19]. The TF-MoDISco approach should capture motifs that are not just recognized but used by the model, as the importance scores are calculated with respect to all other parameters and the model prediction score. We note that we are not the first to widely benchmark importance scores—Prakash et al.[42] evaluated importance scores’ ability to locally highlight motifs within a given sequence across 5 simulated cell-types. We emphasize, however, that our benchmark is different to this previous benchmark in the following ways: (1) We test importance scores across a wider range of neural network architectures, (2) We simulated several magnitudes more genomic grammars, and, most importantly, (3) We score models based on *global* interpretability, in which we evaluate whether a trained model-importance score is capable of learning relevant motifs, rather than their *local* interpretability, where they evaluated whether importance scores were higher for nucleotides in which a motif was implanted compared to background nucleotides. While relevant for local interpretation, we also argue that this metric can at times be mis-calibrated with human-level interpretability—a model may assign high importance to a group of nucleotides that noisily resembles or is close to an actual motif, but where a ground-truth motif has technically not been implanted. By assigning high importance at this region, the model is in fact correctly highlighting a relevant region of interpretable biology, but will be scored as a control under this type of evaluation.

Broadly, causal motif discovery power was quite high (average across all xAI methods, network architectures and simulations settings = 0.723 95% CI[0.713, 0.731]), generally increasing as a function of sample size (AUCs 0.634 [0.622,0.646], 0.742 [0.732, 0.753], 0.793 [0.780, 0.805] at sample sizes of 20,000, 100,000 and 300,000, respectively) and simplicity of genetic architecture (from fully additive to fully epistatic AUCS 0.765 [0.749, 0.780], 0.711 [0.694, 0.726], 0.693 [0.678, 0.707] respectively; Fig. 3).

**Figure 3:**
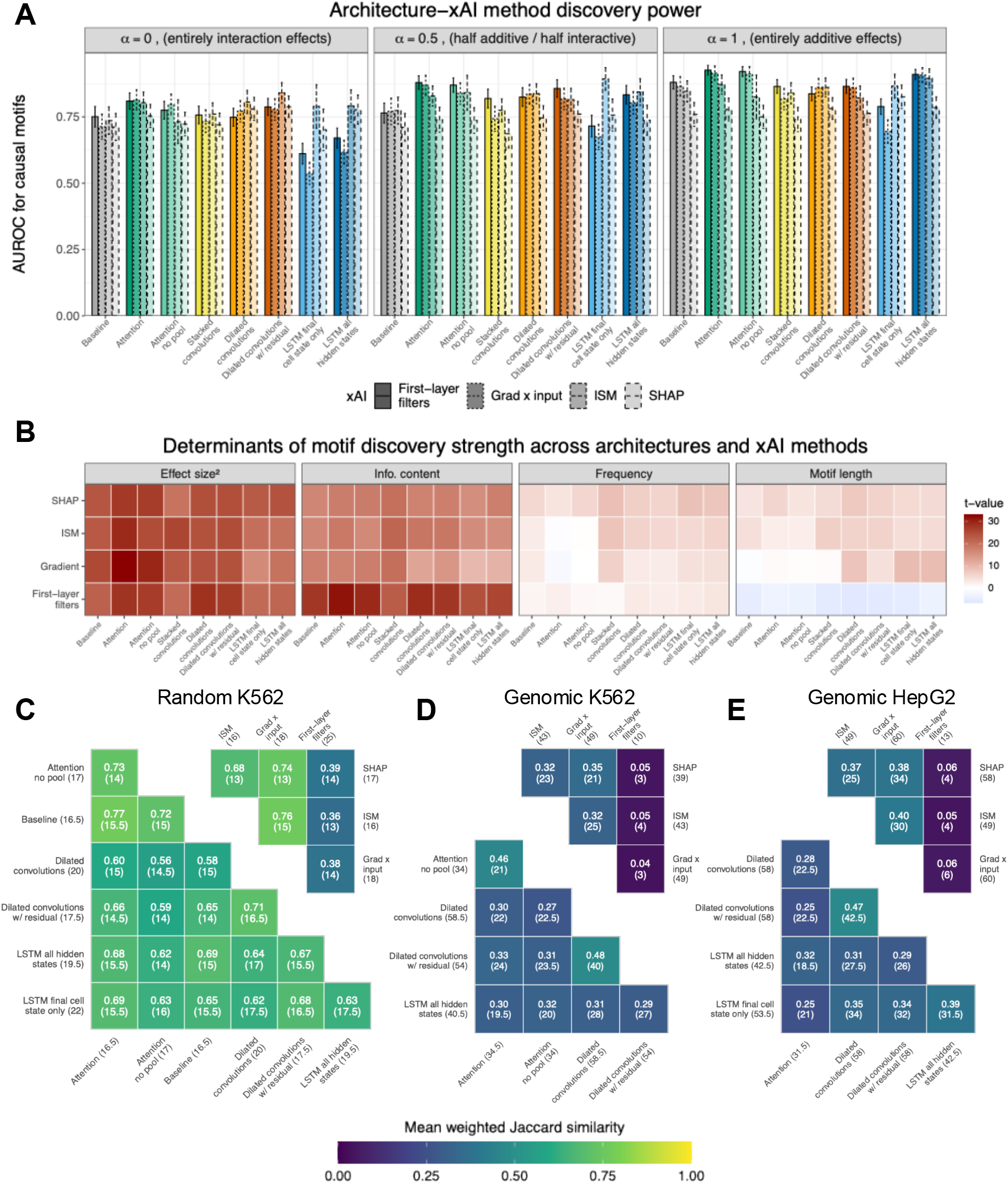
Interpretability methods overlap substantially in motifs discovered. (A) The AUC of each model-xAI pair for prioritizing causal motifs across genetic architectures and under a sample size of 300,000 sequences. Error bars represent 95% confidence intervals obtained via non-parametric bootstrapping. (B) Association statistics between motif characteristics and discovery strength for correctly identified causal motifs across all simulation settings. See Supp. Figs. S3-S8 for individual settings. (C-E) The average weighted Jaccard similarity of motifs discovered within an xAI method (model architecture) across model architectures (xAI method) for (C) the random sequences in K562 cells of Frömel et al. as well as (D) the genomic sequences of Agarwal et al. in K562, and (E) HepG2 cells. Numbers in parentheses indicate the median number of motifs discovered and median number of motifs in common to both xAI method (or architecture) across 10 training runs. Included are architectures whose median prediction accuracy (across 10 training instances) achieved at least 15% of the best-performing architecture.

For the purpose of brevity, we next describe in detail the results from models fit with 300,000 data points in the absence of non-motif effects. Interestingly, simpler approaches (first-layer filters and gradient × input importance scores) were consistently most powerful for the attention-based and baseline approaches (t-test of first-layer filter and grad × input AUC compared to ISM and SHAP within attention and baseline models as a random effect t=3.6, p=9.8e-04). Across all models, first-layer filters were significantly more powerful when the genetic grammar was entirely additive compared to when it was epistatic (association t-statistic of AUC and additivity, accounting for model as a random effect, = 8.17, p=6.67e-07), potentially highlighting models’ necessity to form more complex, hierarchical or composite motifs to effectively model motif interactions (suggested previously in [19]). Within CNNs and LSTMs, *in silico* mutagenesis (ISM) was the most powerful approach in the most difficult genetic grammar (fully epistatic; t-test of AUC as a function of ISM or not while accounting for model as a random effect = 3.02, p=0.009), though, within CNNs, these differences were not statistically significant (biggest difference: dilated CNNs without residual connections and ISM AUC, 95% CI = 0.813 [0.763, 0.862] compared to first-layer filters 0.746 [0.702, 0.790]). Interestingly, SHAP importance scores were underpowered compared to other methods in the strictly additive genetic grammar, likely because they are calculated in reference to a set of background sequences, which, in the absence of context-specific (e.g., epistatic) effects, are effectively independent and do not boost discovery power (t-test of AUC using SHAP vs. all across models as a random effect = −5.20, p=2.86e-05; SHAP vs. ISM and grad × input alone t=-4.16 p=4.1e-04). Across other simulation settings, where sample size was limited, or where motifs had weaker effects (due to non-motif variability affecting the phenotype), differences were generally less significant, and grad × input provided robust discovery power (Supp. Figs.S3-S8).

We next evaluated characteristics of correctly identified causal motifs to understand tendencies or biases toward their discovery across models. In other words, we looked at, among the motifs that had a non-zero effect and were statistically significantly discovered, whether there existed associations between their discovery power and effect size, information content (IC), frequency, or length. Under a specific genetic architecture and across all simulation realizations simultaneously, we fit for each xAI-architecture pair a linear model of motifs’ −log10 discovery p-value onto IC, frequency, squared effect size and length (Methods).

Pooling across models, xAI methods and simulation settings, squared effect size had the largest association statistic to discovery power (absolute mean and 95% CI 22.67 [21.77, 23.62]) followed by information content (18.43 [17.40, 19.51]), motif length (4.69 [4.15, 5.19]) and motif frequency (4.42 [3.68, 5.06]). Nonetheless, at the individual level, IC was significantly more important than effect size for the first-layer filter approach (pooled across all network architectures t=-2.84, p=0.014). Motif length was generally moderately positively associated with discovery power across all xAI methods except for first-layer filters (pooled average association statistic 4.60 [3.94, 5.28] compared to −4.90 [−5.45, −4.34])—likely because (1) a longer motif creates longer contiguous stretches of higher attribution, assisting their discovery and clustering by TF-MoDISco against noisy, sporadic background attribution, signals and (2) any motif that is longer than a specified filter length (here, 7), will be harder to represent in a single filter. This effect was particularly more pronounced for LSTMs and dilated CNNs. Finally, we wished to check for xAI robustness to motif characteristics—in other words, across association statistics, which xAI method produced the smallest associations (and was thus most agnostic to motif characteristics). For all tested architectures, ISM produced the lowest Frobenius norm across models and association statistics (lowest norm of ISM 28.27 compared to next-most agnostic, gradient 31.41 in the strictly epistatic context). Conclusions were largely similar across simulation settings (Supp. Figs. S3-S8).

We then looked at motif discoveries across model-xAI pairs for the three experimental datasets. For each dataset, we examined discoveries for a given xAI-model pair trained across 10 training instances on the full, non-down-sampled dataset, and report for each method the weighted Jaccard index (weighted by number of times each model-xAI pair discovered the motif). In each comparison, Jaccard similarity was not only quite high, but statistically significant (lowest across xAI methods of 0.36, to highest between architectures at 0.77, p*<* 10^*-*3^ permutation test against background motifs). In the random sequences, where each sequence was implanted with several of 44 motifs, the median number of motif discoveries ranged from 16 to 22 across network architectures. Examining genomic-context sequences, our results were largely consistent, however, first-layer filters were substantially less powerful relative to other xAI methods than in the context of random sequences (median 10 discovered motifs, max Jaccard similarity with other xAI methods = 0.05, compared to the random sequences 25 motifs and 0.39 Jaccard similarity). Worth highlighting is the size of the background set of motifs (44 in the random sequences compared to 840 in the JASPAR vertebrate non-redundant database) and thus lower similarities are broadly expected, however, the substantial drop across methods within the genomic-context datasets suggest that first-layer filters are much more robust at distinguishing motifs within a context of random sequences, relative to genomic sequences (where repeated regions likely make the problem harder and filters capture less-informative, non-TF-motif substrings of sequences). Finally, we attempted to validate motif association statistics from our simulated data. Relationships were largely consistent for motif length and information content (Supp. Figs. S9-S11). Nonetheless, for the random sequences in K562, we observed slight negative associations between discovery rate and information content, likely because the regressions conditioned on motif length, but did not include motif effect size—the strongest signal we discovered with the simulated data. This suggested to us confounding by the lack of motif effect size—for which, we emphasize, there is no ground truth—in the association tests. We accordingly turned to current xAI approaches for learning motif effect size via *in silico* experiments.

### *in silico* approaches for interpreting motif effect size rank well but are miscalibrated

To extract model-learned motif effect sizes, we use Global Importance Analysis (GIA), a widely used explainability method to learn feature importance of genomic neural networks[43, 44, 45, 46, 47]. Briefly, GIA, in the simplest case, reports feature “importances”— henceforth used interchangeably with “effect sizes”—by looking at the mean paired difference in model prediction score between a baseline sequence and the same sequence with the feature (motif) implanted in the sequence (Fig. 1). Given we have ground-truth effect sizes for over 1,000 genetic architectures, our analysis represents, to our knowledge, the first large-scale study of GIA faithfulness to the underlying generative model.

We evaluated GIA across varying motif-construction approaches (using the ground-truth PWM to which the model-construction was matched in MEME, the first-layer filter PWM, and the PWM from TF-MoDISco gradient × input report); background sequences (train and test sets); and augmentations (observed sequences, shuffled sequences, and entirely random sequences). When motif grammar is entirely additive, GIA accurately recovers the true motif effect sizes, with the ground-truth PWM motifs (median Spearman correlation [range] across models 0.920 [0.917, 0.940] at N=300,000) slightly outperforming the motifs constructed from the first-layer filters (0.916 [0.785,0.942]) or TF-MoDISco output (N=300,000: 0.886 [0.856,0.918]; Fig 4B, Supp. Figs. S12-S17). In the presence of epistatic effects, the correlation between main effect sizes and their GIA estimates remained consistently high for motifs that did not have an epistatic effect (median [range] for TF-MoDISco-constructed motifs at N=300,000: 0.857 [0.733,0.921]; N=100,000 0.851 [0.710, 0.931]; N=20,000 0.822 [0.671, 0.858]), however, it dropped significantly for motifs involved in epistasis (N=300,000 0.701 [0.614, 0.749]; N=100,000 0.742 [0.621, 0.819]; N=20,000 0.766 [0.676, 0.849]; Supp. Figs. S12-S17). Interestingly, first-order GIA experiments also captured significant variability within of motifs’ mean epistatic effect in the purely epistatic contexts (median [range] correlation between first-order GIA effect and mean epistatic effect across all epistatic contexts 0.601 [0.322, 0.734]), suggesting that it can help prioritize motifs even when the grammar is mis-specified by the *in silico* experiment. We conclude that GIA-estimated importance was highly correlated with the ground-truth main effect sizes, providing evidence for its robust ability to rank motifs over their influence on the modeled outcome.

**Figure 4:**
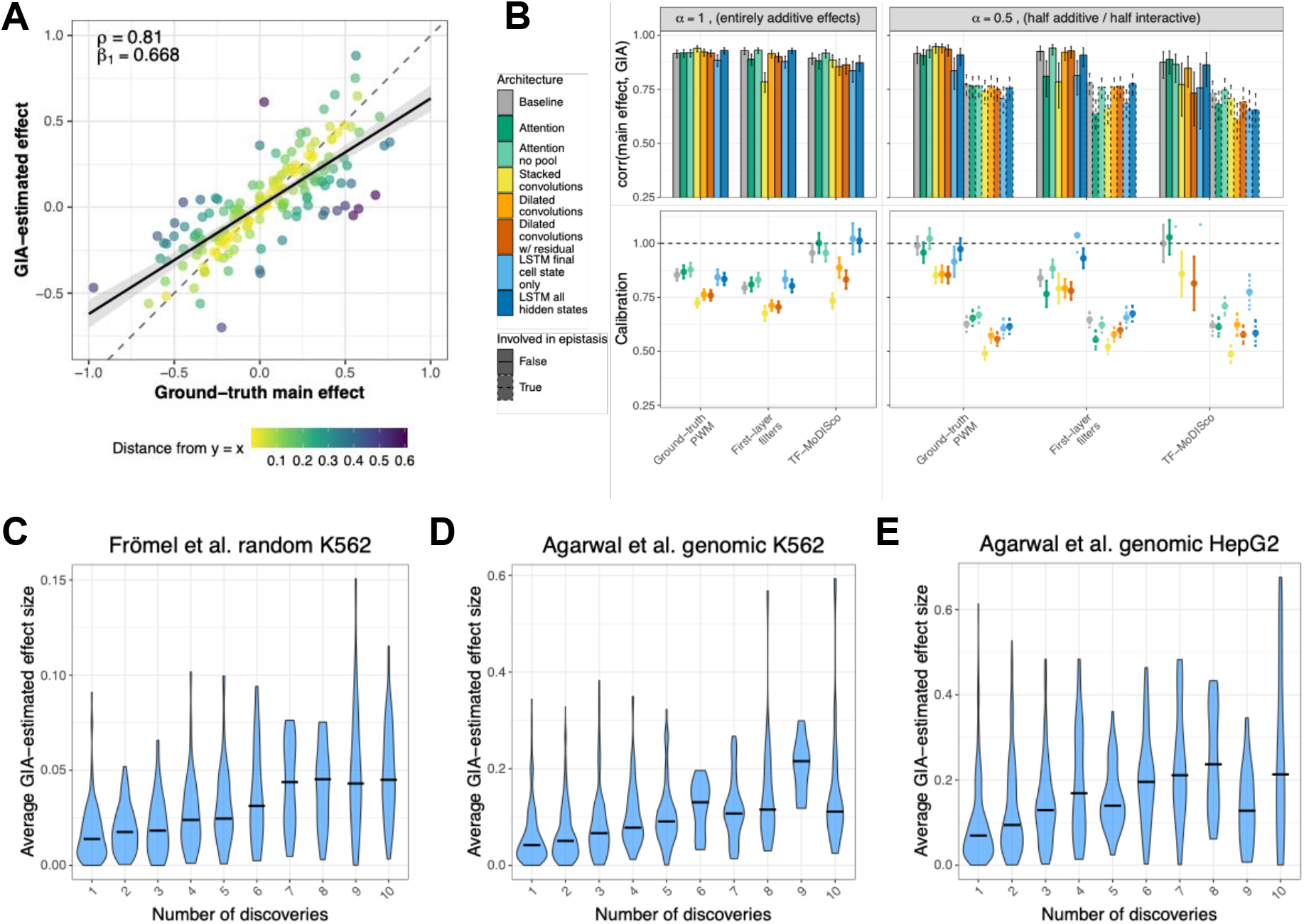
*in silico* approaches highly replicate underlying motif effect sizes. (A) Example correlation and calibration (slope) of ground-truth motif effect size and its corresponding GIA-estimated effect size for a given simulated genomic grammar pooled across 50 realizations. (B) Summary statistics as derived in (A), but across models, genomic grammars, and different xAI strategies for constructing motif PWM for use in GIA. (C-E) GIA effect sizes for real data as a function of number of times each motif was discovered on the Frömel et al. random sequences (C), Agarwal et al. K562 genomic sequences (D), and Agarwal et al. HepG2 genomic sequences (E).

Though our simulated train and test sequences were drawn from the same random uniform distributions, we evaluated differences of GIA-reported effect sizes across train-test splits and data augmentations to evaluate any cases of overfitting or model memorization. Correlation between GIA effects estimated on the simulated train and test sequences was high across models and sample sizes (median Spearman correlation 1.00 [0.944, 1.00]). Subsequently, as expected, using either a set of a random background sequences or the shuffled training sequences as input for GIA led to nearly identical motif effect size estimates (median spearman correlation across models, sample sizes, motif additivity 0.999 range [0.867, 0.999]). The relationship between GIA effects on un-modified training sequences compared to their ablation (via shuffling) was less trivial. For example, in the half-epistatic, half-additive grammar settings, among motifs with only main effects, correlation between GIA-effects estimated on the observed training sequences compared to their shuffled counter-parts remained high (median [range] across sample sizes, models = 0.940 [0.932, 0.970]), whereas if they also had epistatic effects, dropped (0.887 [0.874, 0.952]). In the context of entirely epistatic effects, this correlation worsened (0.518 [0.146, 0.695]). This difference is indicative that the model has learned to contextualize the motif in the training sequences, which we expand upon in the final Results section.

Apart from evaluating ranking of motifs, we also examined how well the GIA-learned effect sizes captured the correct scale of ground-truth motif effect size. To do so, we ran a simple linear regression of the ground-truth effect size onto the estimated importance score, and treated the slope as a metric of calibration (where a slope of 1 indicated the estimate is on the same scale as the true effect size, a slope below 1 indicated over-estimation of effect size, and a slope greater than 1 under-estimation). For motifs in the entirely additive context, all methods and motif-construction techniques were reasonably calibrated (across models at N=300,000 grountruth PWMs median [range] calibration 0.780 [0.724 0.878]; TF-MoDISco PWMs 0.918 [0.733, 1.00]; first-layer filter PWMs 0.754 [0.676, 0.832]). In the half-additive, half-epistatic context, calibration between the GIA estimate and true main effect of motifs involved with epistasis worsened significantly compared to motifs that were not involved with epistasis but also still had a non-zero main effect (median calibration at N=300,000 across PWM strategies and models 0.860 [0.764, 1.02] compared to 0.611 [0.486, 0.721]).

Finally, we examined GIA effects across experimental datasets. We first verified that GIA effect size was indeed associated with motif discovery by regressing the number of times a motifs was discovered per given number of model-training instances in a binomial random effects model that account for network architecture and found highly significant associations for the random K562 sequences of Frömel et al. and both genomic-context sequences of Agarwal et al. (K562, HepG2; Fig 4C-E). Moreover, across datasets and PWM construction strategies, the GIA effects between the shuffled and un-modified sequences remained highly correlated (using ground-truth PWMs across models = 0.981, 0.989, 0.992, TF-MoDISco 0.968, 0.990, 0.987, and first-layer filters 0.976, 0.991, 0.991 for the random K562, genomic K562, and genomic HepG2 cells respectively). Finally, correlations of GIA effect sizes across train and test sequences remained high in the random K562 sequences (median [range] correlation across models 0.9997 [0.9980,0.9999]), K562 genomic sequences (0.9999 [0.9574, 0.9999]), and HepG2 genomic sequences (0.9999 [0.9846, 0.9999]). We conclude that GIA estimates are robust for ranking motif importance across training and test splits, but may be misleading in the context of epistasis.

### Implicating motifs with epistatic effects remains labor-intensive

We next turned our focus to motif interactions, or epistasis. We note that there exist a multitude of ways to extract model-learned motif epistasis, however many of these methods are either local explanations that don’t trivially reveal global genetic grammar (e.g., Deep Feature Interaction Maps [48]), or are quadratic in the number of motifs (e.g., CREME [49]), and therefore computationally prohibitive for our evaluation over 1,500 simulations with our architecture-saliency-motif hyperparameter grid. Accordingly, we examined whether using linear-order approaches (GIA) could implicate epistasis-involved motifs. We hypothesized that while the classic GIA approach of taking the mean perturbation score over many sequences can learn motif main effect sizes by integrating out sequence-specific noise, sequence-specific noise is in fact a desirable quantity for learning higher-order (epistasis) involvement. In brief, large variance in the motif’s perturbation score across sequences may imply that certain backgrounds coordinate with the injected motif, and that this background coordination (or dispersion therein) may be due to an additional motif in the background sequences. Accordingly, we evaluate, in addition to the mean local GIA importance scores across sequences, the variance of local GIA importance scores across sequences as a predictor of motif involvement in epistasis (Methods).

We first examined our metrics in the genetic grammar of half-additive and half-epistatic motif effects. We calculated each GIA metric over the original training sequences, the shuffled training sequences, as well as the pair-wise difference in each (referred to here as “delta”). Intuitively, the delta approach serves as a baseline comparator for learning epistasis or context-specificity as it probes whether the motif effect is specific to the sequences in which is was trained relative to the composition on which it was trained. Nonetheless, neither GIA on the training sequences, GIA on the shuffled training sequences, nor their mean pair-wise difference were sufficient to robustly implicate whether discovered motifs were involved in epistasis across the simulated genetic architecture with half of the heritability explained by simple additive effects and half explained by interactive effects (at N=300,000 using ground-truth GIA, the median [range] AUC across models for GIA importance on the training sequences, the shuffled training sequences, their pair-wise difference were respectively 0.540 [0.500, 0.564], 0.508 [0.463, 0.551], 0.531 [0.454, 0.580]). However, using the variance, rather than the mean of importance scores across sequences, robustly provided modest, but statistically significant separation of epistatic and non-epistatic motifs (at N=300,000 using ground-truth GIA, the median [range] AUC across models for GIA importance on the training sequences, the shuffled training sequences, their pair-wise difference, or covariance were respectively 0.595 [0.512, 0.615], 0.598 [0.550, 0.623], 0.594 [0.499, 0.621], 0.583 [0.551, 0.606]). We also note that the first-moment GIA metric may cause leakage to the second-moment metric (e.g. heteroscedastic effects of the GIA main effect), for which we also evaluated using the variance in importance across observed sequences after having residualized over the corresponding global importance across observed sequences (“Residualized Var(Seq)”; Methods). Nonetheless, even after removing first-moment information, the metric retained the ability to implicate epistatic motifs (at N=300,000 using ground-truth GIA, the median [range] AUC across models 0.613 [0.505, 0.648]), suggesting second-moment metrics indeed harbor relevant epistatic information. Finally, we compared GIA metrics across PWM construction strategies and found that they were most powerful when constructing the GIA motif to the MEME-matched ground-truth PWM, likely due to noise in construction of the PWM from the first-layer filters or TF-mODISco approaches (though, the approach was still significantly powerful for the latter approaches Supp. Figs. S18, S19)

We also evaluated whether specific GIA metrics were more powerful for different types of interactions. To do so, we measured the AUC using the GIA metric as a predictor variable and assigned the classification label as a 0 if the motif was not involved with an interaction and a 1 for a specific interaction type if its largest interaction effect size was for the corresponding interaction type. Interestingly, first-moment GIA estimates showed modest signal for the “simple” interactions (where presence of both motif A and motif B is sufficient for interaction effect) and the “high-order” effects (where presence of all motif A, B and C is necessary for an interaction effect): at N=300,000 using ground-truth GIA, the median AUC across models for GIA effect on the training sequences, the shuffled training sequences, or their pair-wise difference respectively for the simple interactions: 0.624, 0.549, 0.606; or higher-order 0.535, 0.503, 0.551. See Supp. Figs. S20, S21). This was likely due to some non-zero linear dependence in the main effect of the motif presence and the indicator variable representing presence of both (or all three) motifs. In any case, the second-moment GIA metrics were generally over 10% more powerful than the first-moment metrics for both broad separation (at N=300,000 using ground-truth GIA, the median AUC across models for GIA variance on the training sequences, residualized variance, the shuffled training sequences, their pair-wise difference, or their covariance respectively for the simple interactions: 0.658, 0.602, 0.666, 0.689, 0.576; or higher-order 0.597, 0.638, 0.543, 0.614, 0.521. Fig. 5.) and top-1 precision, in which the methods were scored for enrichment of epistatic motifs in their highest values of the GIA metric (GIA on the training sequences mean compared to variance 1.01x to 1.12x enrichment; on the shuffled sequences 0.95x to 1.07x; their pair-wise difference 0.99x to 1.07x; ratio 1.08x enrichment; covariance 1.11x enrichment Fig. 5.). Interestingly, epistatic information for the “upstream” interaction (where motif A must fall upstream of motif B in the sequence for an interaction effect) was more readily extracted when using GIA metrics on the shuffled training sequences, rather than the unperturbed sequences (median AUC across models on N=300,000 sequences, ground-truth GIA using variance on the observed training sequences 0.579, its residuals 0.662, shuffled training sequences 0.654, their delta 0.503, their implied covariance 0.766; Supp. Figs.S20, S21). Finally, we note that nearly all approaches (except for the residualized variance) were under-powered for detecting motifs whose epistasis was distance-dependent (the “distance” interaction, requiring maximum 3 basepairs of distance; median AUC across models using ground-truth GIA at N=300,000 sequences for the importance variance on observed training sequences 0.525, its residuals 0.604, the shuffled training sequences 0.528, their pair-wise difference 0.555, their implied covariance 0.494. Supp. Figs. S20, S21). This is likely due to the fact that when randomly inserting a motif into the sequence, the probability that the motif is injected into a sequence with its partner sequence *and* that the injection happens within the distance threshold (here, 3 nucleotides) is quite low, and hence the variance is modest. This explanation also helps explain the lower power for the high-order epistasis motif detection, as the probability of being injected into a sequence with the other two motifs is at least one magnitude lower than the simple case.

**Figure 5:**
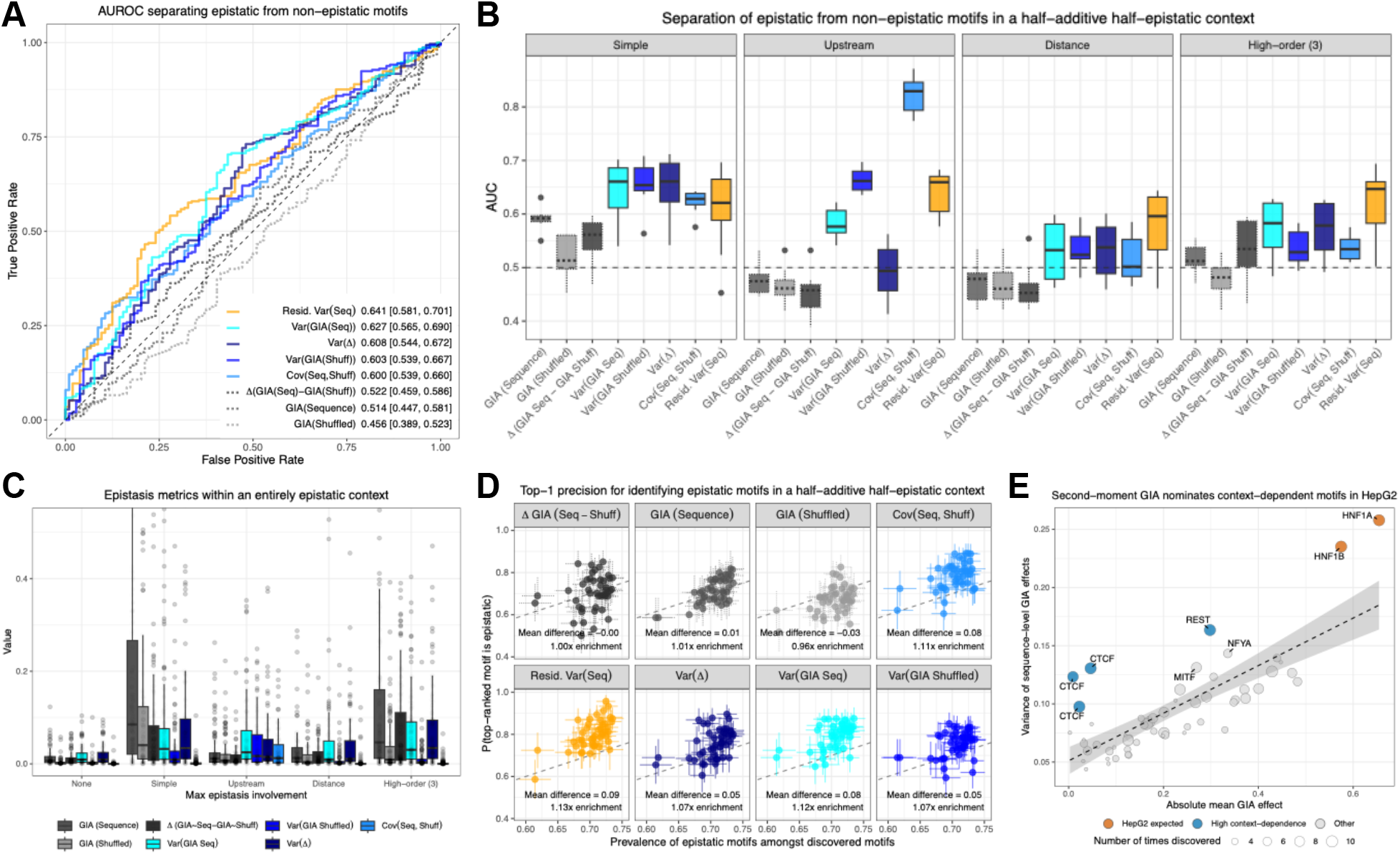
Using first-order *in silico* approaches to implicate epistatic motifs. (A) Distinguishing discovered epistatic from non-epistatic motifs in an example architecture (Dilated Convolutions) with a variety of first-order GIA experiments. Metrics capture variance and mean GIA effects on the original training sequences, the shuffled training sequences, or their pairwise difference—Var(·) implies taking the variance of the importance score, Resid. Var(Seq) represents the variance in local importance on the observed sequences (second-moment) after residualizing over the global importance (first-moment) (B) The distribution of AUCs from panel (A) across each neural network architectures in the context of 300,000 sequences, a half-additive half-epistatic genomic grammar, no non-motif sequence effects, and using ground-truth PWMs for GIA. (C) The distribution of absolute GIA metrics across architectures in the entirely epistatic (absence of first-order main effects) genetic grammar with 300,000 sequences, no non-motif sequence effects, and using ground-truth GIA (See Supp. Figs.S22, S23 for others). The residuals are excluded, as by construction, they center around 0. (D) Scatter-plot quantifying top-1 precision (i.e., how often the top-GIA metric motif is a motif involved in epistasis) as a function of the proportion of discovered motifs involved in epistasis. Error bars represent respective standard errors. Points are shown for all PWM-construction methods and for settings where the phenotypic variability included effects from non-motif features. (E) Applying GIA metrics to the Agarwal et al.[41] genomic-context sequences in HepG2 cells across the 10 model-training instances of the Dilated Convolution network, where each point captures the mean GIA variance as a function of its mean GIA effect across training-discovery instances.

We also applied the above GIA metrics to the 3 real datasets, focusing on motifs enriched for variance in the GIA effect on the training sequences as well as those enriched for implied covariance between the observed and shuffled training sequences as these metrics showed the best discovery power in the simulated data. We begin with an exploration over the K562 random sequences of Frömel et al., given that the authors discussed at length motif effects and their interactions[40], and the study design included a known ground-truth of implanted motifs. Consistent with their report, we found Fli1 to have a large effect size and large inflation of second-moment GIA importance—they found that any motif in the presence of Fli1 will have its effect nullified, effectively leading to an always activated state (Supp. Fig. S24). Further, they reported that having Gata1/2 and Spi1 motifs in background sequences affected readouts of other implanted motifs; across models we frequently found these motifs to have higher epistasis metrics than expected when conditioning on GIA main effect size. Finally, we highlight Mecom, a motif which consistently had one of the lowest GIA effect sizes but highest variance values, suggesting that it would be missed by a typical xAI workflow in the absence of our metric. Interestingly, Mecom was one of the few TF motifs that, in K562 cell-lines, showed both significant activating and repressing activity depending on presence of other background motifs.

Next, we explored motifs implicated in the Agarwal et al. genomic-context sequences. In the HepG2 cell-line, the motifs with the highest second-moment GIA metric were HNF1A and HNF1B, and, while paralogs (occupying a similar sequence space), had different GIA effects and variances (Supp. Fig. S25). Nonetheless, these motifs are known to act as both homodimers of themselves, or heterodimers between each other, and serve as a biological sanity check for the second-moment metric. More interestingly, when we examined motifs whose GIA variance was greater than expected based on their GIA mean effect, we found repeated evidence for CTCF—a known regulator involved in genome organization and assisting of promoter-enhancer contacts (for our purposes, epistasis)—as well as REST, a known repressor. When we examined the K562 cells of Agarwal we found the largest epistasis enrichment for ATF4 and CEBPG—two transcription factors which frequently heterodimerize—as well as CTCF, which again had low average effect size but large variance (Supp. Fig. S26).

## 3 Discussion

Neural networks enable biological discovery owing to the way that they can automatically construct features and use them to predict a variety of mechanisms. Here, we evaluated current abilities to extract some of this learned biology by using a simulations framework in which biological motifs make up most of the variance in a sequence-phenotype relationship, and later validated our findings on real experimental datasets. We found that attention architectures work particularly well in low-data regimes, saliency scores tend to converge to discovering the same sets of motifs, current xAI approaches robustly reveal first-order importance of motifs, and finally, that using second-moment measurements of xAI approaches can help to implicate motifs involved with epistasis.

Importantly, our evaluation has several limitations. Primarily, we focused our simulations on random sequence data, as we believe experimental data (rather than natural sequences) will become increasingly important for elucidating complex genetic grammar[7,8,9,10,40,44,50]. While we validated our findings on genomic-context data, future work could perform a similar simulations-based evaluation using natural sequences as the background. Second, our models and dataset sizes were only roughly ~ 250,000 parameters and at most 300,000 samples measured for short sequences of lengths ranging from 2-300 base-pairs. Experimental data is increasing in availability, and it is common for models to reach many millions of parameters[24]. We attempted to show what happens in different sample-size-to-parameter ratios, but perhaps a worthy next direction is expanding these ratios and scenarios. Though massive experimental datasets comprise sequences that typically fall in the 100-300 base-pair range[7, 10, 40, 50], it may be worthy to study model behavior across larger sequences, when motif signal is sparser. Moreover, we limited our analysis to 8 fixed architectures. Generally, model building may encompass more architecture choices, but almost always includes hyperparameter optimization within a given architecture regime. Of course, further hyperparameter searches may affect the results, but our goal was to discuss the relationship of the generative model and predictive performance as well as to explore the effect of architectural inductive bias on a fixed parameter budget, for which we only probed the effects of pooling, skip connections, and in LSTMs using all sequence or final state information. Nonetheless, by running the evaluation over 1,500 scenarios, and replicating our findings on experimental datasets, we are hopeful that our findings are indeed generalizable.

We believe the work presented here may provide motivation for several future works. Namely, given the miscalibration of first-order motif effects in interactive genomic grammars, it would be useful to probe whether these biases are resolved when doing higher-order xAI or *in silico* analyses. Owing to the massive number of experiments and parameters in our manuscript, we believe that such an effort currently falls out of scope. Next, it would be worthy to study whether large, pre-trained foundational models that are fine-tuned on MPRA experiments replicate findings of small models that are trained on the MPRA experiments alone. Apart from these directions, we believe that the work presented here will be useful to computational biologists wishing to model their sequence-to-function data as well as methodological researchers aiming to increase interpretability or performance of sequence-to-function models.

## Acknowledgments

We would first like to thank Rosa Martinez Corral for initial discussions and push to turn this into a paper, without her motivation this manuscript would not have happened. We appreciate useful discussions with Lars Velten about the Frömel et al. dataset. We also thank Peter Koo for his helpful comments as well as Anshul Kundaje and Dmitry Penzar for their online suggestions. Finally, we are grateful to the UCLA hoffman2 server and support.

## Author contributions

Conceptualization: M.T. Computational analyses: M.T. Software: M.T. Manuscript writing: M.T. with input from B.L. Visualization: M.T. Funding acquisition: B.L. and M.T.

## Methods

### Simulations

To evaluate neural network architectures and motif discovery methods, we simulated ground truth data with a variety of tunable parameters. Importantly, we set our basis of simulations upon random sequences. Random sequences enable researchers to overcome problems of overly optimistic predictive performance due to homology, as well as to learn more generalizable sequence and motif grammars, since random sequences comprise a much wider range of contexts than the genome[51]. Additionally, as researchers continue to perform experiments using synthetic sequences and random sequences[7,8,9,10,40,44,50], it is crucial to understand the circumstances in which neural networks produce predictions that are faithful to the underlying biology in order to maximize the insight gained from these experiments.

Accordingly, the background sequences comprised 200 base-pairs with a uniform distribution of nucleotides at each position. Into these sequences, we injected up to 10 motifs downloaded from JASPAR[52], each with their own probability drawn from a Uniform distribution (half of the motifs came from Unif[0.1, 0.4] and the other half Unif[0.4, 0.8], distributions were shuffled per simulation run). We preprocessed motifs from JASPAR to create a set of 5 length-7 motifs, as well as 5 others between lengths 4-12 and across a variable range of information content. To increase motif variability, the motifs per sequence were injected as a realization from the collected position-weight matrices (PWM). After drawing and potentially injecting motifs into a sequence (into non-overlapping positions drawn from Unif[1, 200 − |motif|] per motif and sequence), a phenotype was generated following a basic variance components model:

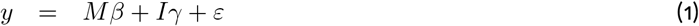

Where *y* is a length-*n* vector of the phenotypes across the *n* sequences, *M* is a scaled *n* × *p* matrix of sequences’ presence of *p* motifs, *β*is length-*p* vector representing the effects of the motifs, *I* is a scaled *n* × *q* matrix across sequences representing the presence of multiple motifs in the arrangement required for them to interact (specified below), *γ* is a length-*q* vector representing the interaction effects, and *ε* is a length-*n* vector of normally distributed noise. Importantly, we define a quantity *h*^2^ which we term “motif-explainability,” that represents the variance in phenotype that can be explained by motifs and their interactions. This quantity is analogous to heritability in the statistical genetics field[53]. We partition the motif-explainability into one component of simple additive effects *β*, which act on *M*, and another that is explained by motif interactions *γ*, which act on *I*. To introduce sparsity of motif effects, we sample per simulation run a subset of *π* proportion of motifs to have an additive effect. We split the proportion of motif-explainability across motifs and their interactions via a parameter *α*, which when set to 0 generates phenotypes whose variability is explained exclusively by interactive effects, and only additive effects when set to 1. More rigorously, we simulate motif effects *β* as:

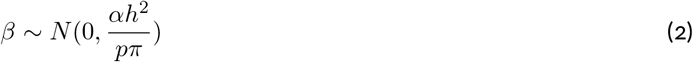

interaction effects as:

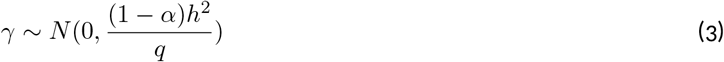

and finally draw random noise *ε* from:

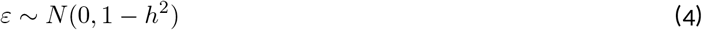

For the purpose of simplifying the simulations, we limit our evaluations to situations in which the motif explainability is set to 80%, there are only 10 motifs (6 of which have an additive effect) and a set of 4 interaction classes among at most 3 hypothetical motifs A, B and C (which are randomly selected from all 10 motifs per interaction and per simulation run) as follows:

#### Simple interactions

The presence of both motif A and motif B at any position in the sequence is sufficient for them to interact.

#### Order interactions

Motifs A and B only interact if motif A is upstream of motif B.

#### Distance interactions

Motifs A and B only interact if the distance between them is less than 4 basepairs.

#### High-order interactions

The presence of motifs A, B, and C produces an interaction.

In the text, a given simulation run refers to a specific realization of background sequences, motif frequencies, motif presence, motif positions, motifs with non-zero effect sizes, sets of motifs involved in the interactions, beta and gamma effect sizes, and noise all drawn under one specific combination of *α, ψ*, and sample size parameters. We performed 50 simulation runs for each distinct combination of parameters. We also evaluated scenarios in which there are non-motif, sequence-based effects on the phenotype (Appendix). We include the simulations script, Probabilistic Evaluation of genomic NEUral networks (PRENEU) and all accompanying code under https://github.com/mj-thompson/PRENEU.

#### Defining a simulation run

Importantly, we present data from a variety of simulation-parameter settings. For each combination of parameters (the sample size as well as the amount of variability due to additive motif effects, interactive motif effects, and the sequence-level effect), we generated 50 realized datasets, then trained each model one time on these 50 realized datasets. One data realization of the simulations consists of the following steps:

1. Generate length-200 random sequences
2. Draw 10 motif frequencies, 5 from Uniform[0.1, 0.4] and another 5 from Uniform[0.4, 0.8]. Shuffle them, so that any motif can have a frequency from either distribution.
3. Using these motif frequencies, sample for each sequence a motif presence indicator variable from Bernoulli(motif frequency)
4. For each sequence, sample a potential location for each motif from Uniform[1, 200 − length(motif)]. If any motif overlaps, resample motif positions until they do not.
5. For each random sequence, realize a motif sequence from its corresponding position weight matrix and overwrite the random sequence (starting from the motif’s realized position to the motif’s length) if the motif’s corresponding realized Bernoulli random variable was 1
6. Calculate each sequence’s non-motif effect, if present (below)
7. 7. Randomly sample 3 pairs from the set of 45 (10 choose 2) pairs of pairwise interactions for the interactive effects. Also sample 1 trio from the set of 120 (10 choose 3) trios for the higher-order interactive effect.
8. Sample *(β, γ*, and *ε*, then use them with the calculated non-motif effect to assign a phenotype for each sequence.

#### Calculating non-motif sequence-level effects

In addition to the motif-based effects on the phenotype, we considered non-motif effects, inspired by alternative splicing. The idea was to quantify how “intron-like” the sequence is. In other words, we further complicated the problem by introducing a non-motif effect *S* whose contribution to the phenotypic variance depends on *ψ*. This effect searches for AG dinucleotide enrichment at the start of the sequence and polypyrimidine enrichment at the end of the sequence, then calculates a potential *S* based on the distances between the enrichments (Appendix). The effect can be reified as an “alternative splicing-like” effect that prefers sequence characteristics and a specific distance between them, though more importantly, the focus of the effect is to increase difficulty of motif-discovery. In such cases, the phenotype is specified as follows:

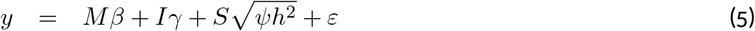

and *(β* and *γ* are sampled as before, only with an additional scale of 1 *-* in the numerator of the variance.

The scoring mechanism first scans left-to-right, quantifying the “intron-potential” of a sequence by scoring for the presence of a downstream polypyrimidine tract. The potential rises (and eventually fires) in a stretch of polypyrimidines, and declines in the presence of non-polypyrimidines. After spiking, there is a refractory period. In other words, this polypyrimidine intron potential looks for an intron-exon border via the polypyrimidine tract, and after finding a possible intron-exon border enters a cool-down state, where no new possible border can be characterized. The right-to-left score works similarly, only that it traverses the sequence in the opposite direction while scanning for enrichment of the canonical exon-intron dinucleotide splice site motif of “GA.” The final score is calculated by combining the left-to-right and right-to-left scores, then penalizing (or rewarding) the scores if the potential bursts occurred close to the ends of the sequence (i.e., there is a sufficient minimum distance between the proposed exon boundaries, implying a reward for reaching at least some minimum intron length).

Explicitly, we scan the sequence characters *c*_1_, …, *c*_*L*_ (where *L* is the sequence length of 200), maintaining at each position *i*:

- state_*i*_, the current position’s “intron likeness”,
- adaptation_*i*_, the tolerance after marking a “splice boundary”,
- refractory_*i*_, a brief “exon-mode cooldown” before the next possible intron-exon (or exon-intron in right-to-left) boundary can fire,
- spikeSum_*i*_, the current accumulated boundary potential,
- *t*_*i*_, the index of the most recent putative boundary (“last splice”).

At each position *i* we add to the state, or intron likeness, a proposed *δ*_*i*_, which, for the left-to-right score, is computed as a polypyrimidine reward:

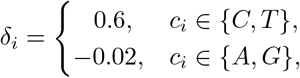

For the right-to-left score, we follow the same logic, only also considering the adjacent position in the sequence:

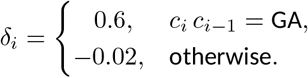

If adding delta_*i*_ to state_*i*_ passes the specified “intron-like” threshold, we then add 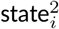 to spikeSum_*i*_ (if the most recent putative boundary occurred at least 10 bases away), update *t*_*i*_ to the current position, reset the state to −1, enter a refractory period of at minimum 5 steps, then increase adaption_*i*_. We then continue to add to the down-(or up-)stream scores the proposed *δ* values, however, we now also consider the refractory period and adaptation (which effectively limit the speed at which a boundary can be called again). In more detail, If state_*i*_ ≥ *threshold* we fire a boundary burst:

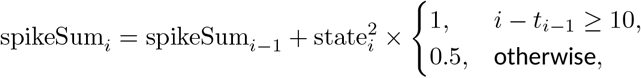

and then

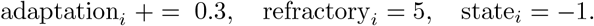

If no burst occurs, spikeSum_*i*_ and *t*_*i*_ simply carry forward.

Once a burst has occurred, then the proposed *δ*_*i*_ to be added to state_*i*_ as well as adaptation_*i*_ are modified as follows:

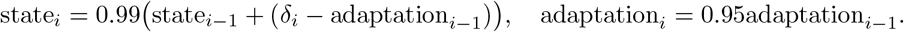

The left-to-right intron-like score then given as:

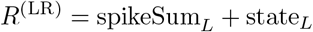

The left-to-right intron-like score *R*^(LR)^ is calculated as specified above for the right-to-left score—in the implementation we simply invert the string and map the string indices to their proper coordinates. In addition to the final scores, we also save for each sequence the location of where their final left-to-right and right-to-left putative boundaries were called, *t*^(LR)^ and *t*^(RL)^ respectively.

Finally, prior to combining *R*^(LR)^ and *R*^(RL)^, we award bonuses when their final proposed features occurred in their expected 5’ and 3’ regions:

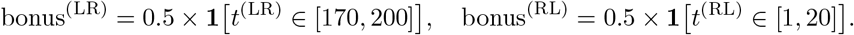

We also apply a penalty if the sequence features occurred too close together, preferring that there is at least 160 nucleotides in between them:

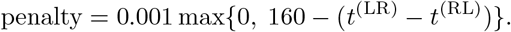

The final sequence-level effect (i.e., intron-likeness score) is then

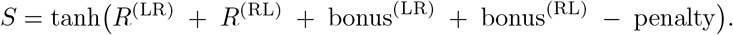

We show below an example of the “intron-like” scores annotated along the sequences below.

**Figure A1:**
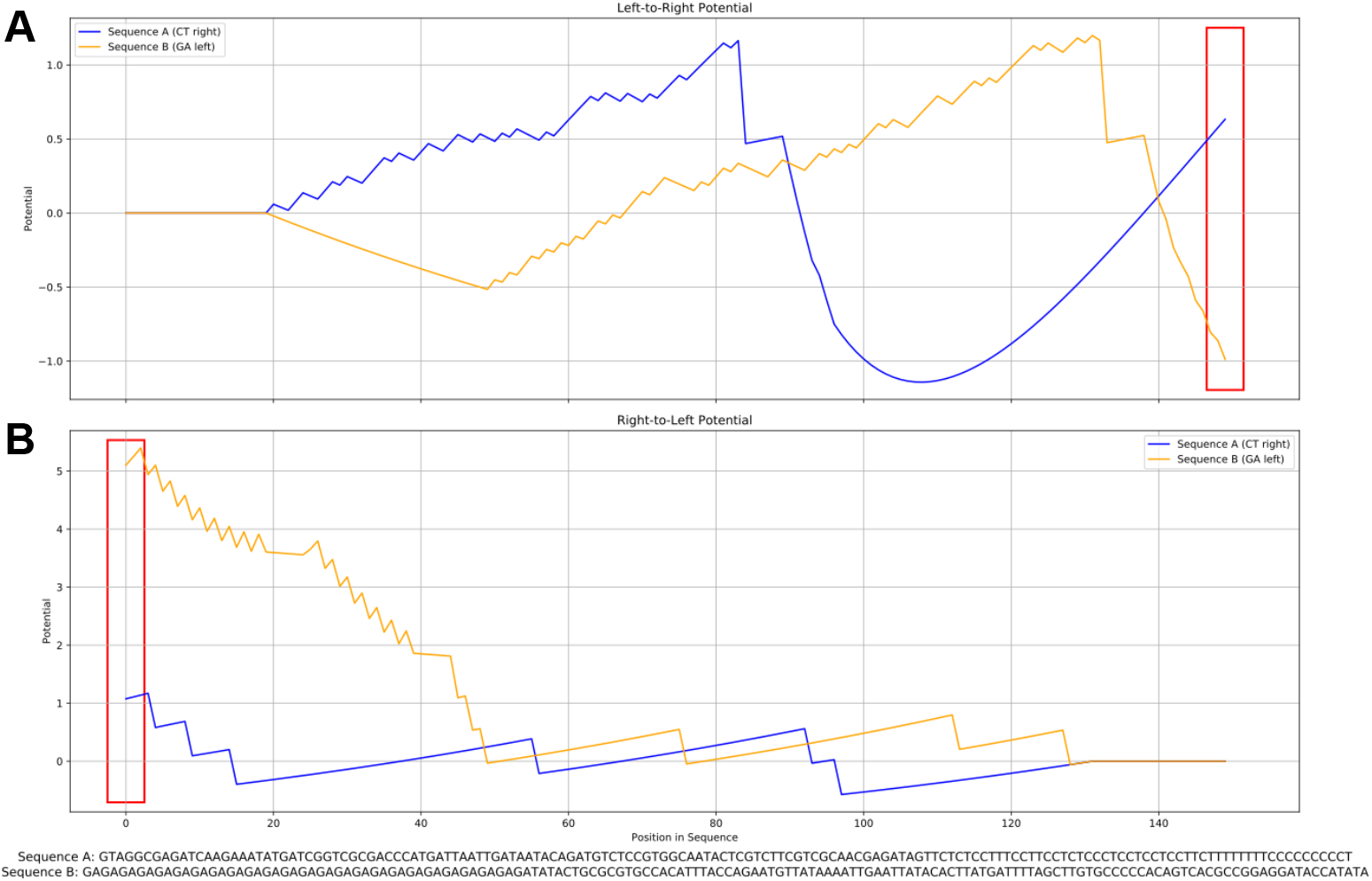
Non-motif sequence level effects. (A) The left-to-right sequence score the searches for downstream polypyrimidine enrichment. The red box represents the final state used in scoring, (*R*^(LR)^).(A) The right-to-left sequence score the searches for upstream GA motif enrichment. The red box represents the final state used in scoring, (*R*^(RL)^)

### Model architecture and training details

In this section, we explicitly define the training routine as well as the layers and hyperparameters (e.g. dropout rate) for all models used in the analysis. All models followed the same training routine, which used 100 as the batch size, 100 as the number of epochs, 0.10 (of the training data) as the validation split, mean squared error (MSE) as the loss, the tensorflow default adam as the optimizer, and finally, early stopping and learning rate reduction as the callbacks. Both callbacks monitored the validation loss and were run in mode “min”. The early stopping callback employed a patience of 10, whereas the reduce learning rate callback used a patience of 6, a factor of 0.2, and a minimum learning rate of 1e-05.

#### Baseline

1. Input layer (size 200×4)
2. 1D convolution layer with 100 length-7 filters, followed by exponential activation, and dropout at rate 0.1.
3. Dense layer with 48 nodes, elastic net regularization of 1e-04, batch normalization, ReLU activation, and dropout at rate 0.4.
4. Flatten
5. Dense layer with 24 nodes, elastic net regularization of 1e-04, batch normalization, ReLU activation, and dropout at rate 0.4.
6. Output layer with 1 node and linear activation

#### Attention

1. Input layer (size 200×4)
2. 1D convolution layer with 100 length-7 filters, followed by exponential activation, max pooling with window size 3, and dropout at rate 0.1.
3. Multi-head attention with 8 attention heads, key length 6, multiple positional embeddings (relative shift, gamma distribution, exponential decay, central-window masking) and dropout at rate 0.10.
4. Flatten
5. Dense layer with 64 nodes, elastic net regularization of 1e-04, batch normalization, ReLU activation, and dropout at rate 0.4.
6. Output layer with 1 node and linear activation

#### Attention no pool

1. Input layer (size 200×4)
2. 1D convolution layer with 100 length-7 filters, followed by exponential activation and dropout at rate 0.1.
3. Multi-head attention with 4 attention heads, key length 6, multiple positional embeddings (relative shift, gamma distribution, exponential decay, central-window masking) and dropout at rate 0.10.
4. Flatten
5. Dense layer with 48 nodes, elastic net regularization of 1e-04, batch normalization, ReLU activation, and dropout at rate 0.4.
6. Output layer with 1 node and linear activation

#### Stacked convolutions

1. Input layer (size 200×4)
2. 1D convolution layer with 100 length-7 filters, followed by exponential activation, and max pooling with window size 3.
3. 1D convolution layer with 30 length-3 filters and ReLU activation.
4. 1D convolution layer with 30 length-5 filters, followed by ReLU activation and max pooling with window size 2.
5. 1D convolution layer with 60 length-2 filters and ReLU activation.
6. Flatten
7. 7. Dense layer with 96 nodes, elastic net regularization of 1e-04, batch normalization, ReLU activation, and dropout at rate 0.4.
8. Dense layer with 64 nodes, elastic net regularization of 1e-04, batch normalization, ReLU activation, and dropout at rate 0.4.
9. Output layer with 1 node and linear activation

#### Dilated convolutions

1. Input layer (size 200×4)
2. 1D convolution layer with 100 length-7 filters, followed by exponential activation, and dropout at rate 0.1.
3. Dilated convolution block consisting of a 1D convolution layer with 100 length-7 filters, ReLU activation, and dropout at rate 0.1, followed by a second 1D convolution layer with 100 length-7 filters, ReLU activation and dropout at rate 0.1
4. Dilated convolution block consisting of a 1D convolution layer with 50 length-7 filters, dilation rate 3, ReLU activation, and dropout at rate 0.1, followed by a second 1D convolution layer with 50 length-7 filters, dilation rate 3, ReLU activation and dropout at rate 0.1
5. Dilated convolution block consisting of a 1D convolution layer with 33 length-7 filters, dilation rate 9, ReLU activation, and dropout at rate 0.1, followed by a second 1D convolution layer with 33 length-7 filters, dilation rate 9, ReLU activation and dropout at rate 0.1
6. Dilated convolution block consisting of a 1D convolution layer with 25 length-7 filters, dilation rate 27, ReLU activation, and dropout at rate 0.1, followed by a second 1D convolution layer with 25 length-7 filters, dilation rate 27, ReLU activation and dropout at rate 0.1
7. Dilated convolution block consisting of a 1D convolution layer with 20 length-7 filters, dilation rate 81, ReLU activation, and dropout at rate 0.1, followed by a second 1D convolution layer with 20 length-7 filters, dilation rate 81, ReLU activation and dropout at rate 0.1
8. Flatten
9. Dense layer with 16 nodes, elastic net regularization of 1e-04, batch normalization, ReLU activation, and dropout at rate 0.4.
10. Output layer with 1 node and linear activation

#### Dilated convolutions w/ residual

1. Input layer (size 200×4)
2. 1D convolution layer with 100 length-7 filters, followed by exponential activation, and dropout at rate 0.1.
3. Dilated convolution block consisting of a 1D convolution layer with 100 length-7 filters, ReLU activation, and dropout at rate 0.1, followed by a second 1D convolution layer with 100 length-7 filters, addition of block input, and ReLU activation
4. Dilated convolution block consisting of a 1D convolution layer with 50 length-7 filters, dilation rate 3, ReLU activation, and dropout at rate 0.1, followed by a second 1D convolution layer with 50 length-7 filters, dilation rate 3, addition of block input, and ReLU activation
5. Dilated convolution block consisting of a 1D convolution layer with 33 length-7 filters, dilation rate 9, ReLU activation, and dropout at rate 0.1, followed by a second 1D convolution layer with 33 length-7 filters, dilation rate 9, addition of block input, and ReLU activation
6. Dilated convolution block consisting of a 1D convolution layer with 25 length-7 filters, dilation rate 27, ReLU activation, and dropout at rate 0.1, followed by a second 1D convolution layer with 25 length-7 filters, dilation rate 27, addition of block input, and ReLU activation
7. Dilated convolution block consisting of a 1D convolution layer with 20 length-7 filters, dilation rate 81, ReLU activation, and dropout at rate 0.1, followed by a second 1D convolution layer with 20 length-7 filters, dilation rate 81, addition of block input, and ReLU activation
8. Flatten
9. Dense layer with 16 nodes, elastic net regularization of 1e-04, batch normalization, ReLU activation, and dropout at rate 0.4.
10. Output layer with 1 node and linear activation

#### LSTM final cell state

1. Input layer (size 200×4)
2. 1D convolution layer with 100 length-7 filters, followed by exponential activation, and dropout at rate 0.1.
3. Bidirectional LSTM with 64 features, dropout rate of 0.2, recurrent dropout rate of 0.2
4. Bidirectional LSTM with 64 features, dropout rate of 0.2, recurrent dropout rate of 0.2
5. Dense layer with 128 nodes, elastic net regularization of 1e-04, batch normalization, ReLU activation, and dropout at rate 0.4.
6. Output layer with 1 node and linear activation

#### LSTM all hidden states

1. Input layer (size 200×4)
2. 1D convolution layer with 100 length-7 filters, followed by exponential activation, and dropout at rate 0.1.
3. Bidirectional LSTM with 24 features, dropout rate of 0.2, recurrent dropout rate of 0.2
4. Bidirectional LSTM with 24 features, dropout rate of 0.2, recurrent dropout rate of 0.2
5. Flatten
6. Dense layer with 24 nodes, elastic net regularization of 1e-04, batch normalization, ReLU activation, and dropout at rate 0.4.
7. Output layer with 1 node and linear activation

### Evaluation on experimental datasets

To validate our findings, we turned to the experimental data from Frömel et al.[40], where the authors generated a collection of synthetic random sequences embedded with a variety of transcription factor motifs and measured their activity in K562 cell-lines (N=57,342, length=256; Methods)[40], as well as from Agarwal et al. [41], where the authors measured transcriptional activity of short, endogenous, genomic sequences in K562 and HepG2 cell-lines. (N=196,664 and 122,926 respectively, length=230) Specifically, we downloaded the Frömel et al. data from their figshare repository at DOI: 10.6084/m9.figshare.25713519.v3 and accessed the training datasets of libraries A, B, C and their controls for the K562 cell line, randomly splitting all sequences into one 90% train-validation set, and another into a test set. For the Agarwal et al. data, we accessed their Zenodo repository at https://doi.org/10.1101/2023.03.05.531189 [54] and partitioned the 90% train-validation and 10% test sets using the sequence base ID, such that a sequence and its reverse complement were both contained in either the train-validation or test set. For a given architecture, we trained 10 instances of the model over the training sequences and report the performance and motif discovery results as calculated over the test sequences. In experiments where we down-sampled sequences, we set 3 seeds and randomly sampled a given proportion of data 3 times, training 10 models for each 3 subsets, for a total of 30 training instances per model-subset pair. For *in silico* experiments (below), we followed the same settings as those used in the simulations, only for the Frömel et al. dataset, we used their collection of 44 reported motifs as the background set, and for the Agarwal et al. datasets, we used the core, non-redundant vertebrate set of 841 JASPAR motifs from 2022 as reported in their manuscript[41].

### Motif scoring and significance testing

#### Generating and scoring position weight matrices from first-layer filters (shallow approach)

There are a variety of ways to generate a PWM from first-layer filters. Primarily they entail choosing a set of sequences on which to calculate filter activation energies (e.g. test-set sequences), then subsetting these sequences to those that activate the filter over some pre-specified activation-energy threshold (e.g. 50% of the maximum activation energy). Nonetheless, using only sequences observed in the data (whether training or test sets) may lead to noisy PWMs, as this approach assumes that the filter’s best-activating sequences appear in the data (which may not be the case if the model and its filters were optimized over a train set and then visualized with an independent test set). Accordingly, we follow the approach taken in [7]. In this text, the authors collect all *k*-mers, where *k* is the filter length, calculate activation energies for these *k*-mers, then finally apply thresholding over the set of *k*-mers to decide which to include in their PWM construction. In our evaluation, we included a one-hot encoded *k*-mer (scaled by its activation energy, after passing through the exponential function) in the PWM construction if its activation energy was over 90% of the maximum activating *k*-mer’s activation energy. To avoid strong reduction in information content, we used at most 10 sequences in the PWM construction.

We then saved all PWMs to a single meme-format file with uniform background frequencies (0.25) for each nucleotide[55]. Next, we ran the tomtom motif comparison tool[55] over these PWMs and the groundtruth motifs with parameters “–verbosity 1”, “-text”, “-dist pearson”, “-norc”, “-thresh 1.” We called discoveries statistically significant if the reported PWM-ground-truth motif association q-value was *<* 0.05.

#### Generating and scoring motifs with TF-MoDISco

TF-MoDISco generates its list of motifs by scanning over a list of sequences and finding subsequences with enriched (non-zero) importance scores. These subsequences can be derived from training or test sets of sequences (or more clever combinations), and the importance scores can be specified via the calculated gradient of the trained model output with respect to the sequence or more involved methods, like SHAP[26] or DeepLIFT[27]. In this report, we used the following scores calculated over sequences in the test set (the heldout 10% of the data): (1) the gradient with respect to model output (i.e. saliency maps), corrected by projecting it onto the simplex[32]; (2) SHAP, using the training sequences as background sequences and the gradient approximation method for computational efficiency; and (3) *in silico* mutagenesis (ISM), where, for each sequence, we calculated its importance score at a given position × nucleotide as the change in model prediction from the wildtype sequence to the corresponding single mutant nucleotide, where each mutant kept the rest of the wildtype sequence constant, and the wildtype nucleotide received the average of these 3 scores at the corresponding position. To generate motifs, we passed the test sequences and their corresponding importance sequences as input to TF-MoDISco and ran TF-MoDISco (Lite implementation) with the following parameters “-s TESTSEQUENCES”, “-a MODEL_SCORE”, “-n 50000”, “-w SEQUENCE_LENGTH”, as we wished to increase resolution of TF-MoDISco by increasing the number of seqlets used and include the whole sequence in the search space. We generated a report and obtained tomtom q-values by running the report command over the output of the motif generation command and with the “-m” flag set to the same groundtruth meme file as the shallow approach tomtom command. We again called discoveries statistically significant if the reported PWM-ground-truth motif association q-value was *<* 0.05.

#### Calculation of motif discovery power and associations

After extracting learned motifs using the first-layer filters or TF-MoDISco approaches, we examined characteristics underlying motif discovery power. In the simulated data, we first calculated each model-xAI pair’s ability to discover a casual motif by evaluating the area under receiver operating characteristic curve (AUROC; with R package pROC::ci.auc) where the label was set as 1 if the motif was causal (had a non-zero effect, either main or epistatic) and 0 otherwise, and the predictor was the xAI method-architecture’s −log10 p-value as reported by MEME.

To characterize determinants of motif discovery strength, we ran regression models that included network architecture, xAI, and discovered motif traits. Specifically, among true positive, discovered causal motifs, we ran for each xAI-architecture pair a regression model of the −log10 p-value onto the true squared motif effect size (set to the squared maximum main effect or epistatic effect), motif information content, motif frequency (which varied across simulation realizations), and motif length, where each variable was centered to mean 0 and variance 1. On the real, experimental datasets, we had 10 trained model instances (and thus chances for motif discovery) and did not have access to ground-truth effect sizes, so we instead ran simple binomial regressions where the outcome was the number of times the motif was discovered (at q *<* 0.05) out of the number of trials (10) modeled over either motif length, information content, or later squared GIA-estimated effect size (see next sections). We examined each characteristic independently when modeling the experimental data as we noticed ground-truth motif effect size was the greatest driver of motif discovery, and hence its absence could confound multiple regression analyses.

We also characterized similarity between sets of discovered motifs for each xAI-architecture pair on the experimental datasets. To do so, we calculated the weighted Jaccard similarity (where weights were the counts, out of 10, of model-training instances where the motif was discovered) of motifs between xAI-architecture pairs, where the background for both sets was treated as the known, implanted motifs in the Frömel et al. dataset, or the JASPAR 2022 core, non-redundant vertebrate motifs’ gene names (as multiple motifs map to a single gene) for the Agarwal et al. datasets. To calculate significance of Jaccard similarities, we used a permutation test with 1,000 randomly sampled sets from the background motifs with cardinality and weights equivalent to the xAI-architecture’s true discovered set. We limited analyses to models whose median prediction performance fell within 15% of the best-performing model for that dataset.

### Global importance analysis experiments of motif main effect size

Global importance analysis (GIA) is an interpretability method used to quantify feature importance (here, motifs) in genomic neural networks[43]. Briefly, GIA estimates the global (across sequences) importance of a feature of interest by measuring the mean pairwise difference in model output between sequences with and without the implanted feature and can be approximated as follows:

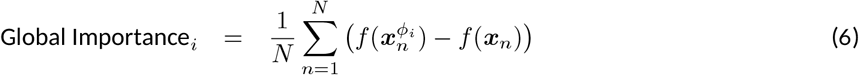

where *N* is the number of sequences used in the calculation, *f* (·) represents the output of the trained model for a given input, ***x***_*n*_ represents the non-augmented sequence 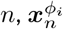 represents the sequence where feature of interest *i* has been implanted, and each term in the summation represents a local importance that captures how the feature affects prediction for each sequence. Technically, the global importance (across all sequences), could be estimated by moving feature *i* along the entirety of each sequence, however, for the purposes of this manuscript, we simply implant the feature of interest along the sequence at a uniformly random position for all sequences, using a large *N* (e.g. size of the entire training or test set) such that most of the distribution of possible position-specific effects is covered. Moreover, for evaluating the importance of motif *i*, we randomly drew a realization of the motif from its PWM (with probability proportional to that specified by its PWM), such that different sequences could contain different realizations of the motif.

For each discovered motif, we ran GIA using either the train or test sequences, keeping the sequences as observed or randomly shuffling them to maintain any composition-specific effects. Additionally, we evaluated the effect of PWM-choice on GIA effect size estimation for given motifs. In other words, for each MEME-discovered motif, we ran GIA when sampling from the ground-truth matched PWM that was input into MEME, the first-layer filter that discovered the motif, or the contribution weight matrix as specified by TF-MoDISco. For the purposes of brevity, we used strict thresholding on *k*-mers used as input to the PWMs for the first-layer filters of TF-MoDISco: in the case of first-layer filters, we chose either *k*-mers with filter activation energy at least 90% of the maximum-activating sequence, using at most the top-10 activating sequences, and in the case of TF-MoDISco, we chose for a motif the implicated pattern with the largest number of seqlets matched to it, trimmed low-information content flanks (<0.2), and again used the top-10 most-attributing trimmed seqlets to align, then build the PWM. As TF-MoDISco can report reverse complements of the matched motif, we also scored via continuous Jaccard similarity the implicated seqlets and their reverse complement, and chose the set with greater similarity as the set for PWM-construction. Here, we used the motifs discovered by TF-MoDISco with gradient (× input) importance scores for run-time’s sake, but note that the PWM is unlikely to drastically change whether using SHAP or ISM, as we use only the top 10 attributing seqlets, which are largely conserved across importance scores.

After running GIA for each model, we examined the relation of GIA-estimated importance to the ground-truth motif effect size using both Spearman correlation between the two, and linear regression with the ground-truth effect size as the outcome. The Spearman correlation enabled us to answer whether estimated importance correctly ranks motifs, and the slope from the simple regression enabled us to evaluate whether the reported global importances were on the same scale (i.e. calibrated) as the ground-truth effect sizes.

### Epistatic motif analyses and second-order statistics

In addition to running GIA as in equation 6, we also evaluated several modifications to the calculated importance to highlight motif involvement with epistasis. Namely, we first examined whether supplying to the mean (a first-moment estimate) background sequence information could help elucidate context-specific effects. In other words, we wished to evaluate whether changes in motif importance on the observed sequence compared to a shuffled version of the sequence could highlight other contexts (i.e. in the presence of other motifs) in which the motif has an altered effect. To do so, we examined the pairwise difference in importance between the sequence and its shuffled sequence, referred to in the main text as “GIA delta”:

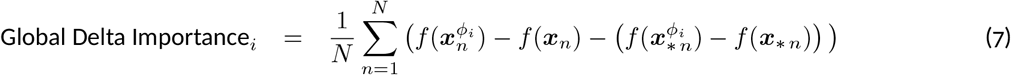

where symbols follow as before and ***x***_∗ *n*_ represents the shuffled sequence ***x***_*n*_, maintaining its sequence composition and both motif-implanted sequences have the same realization from the PWM injected at the same position in the shuffled and unshuffled sequence. Intuitively, the averaging in the original global importance equation effectively integrates over a given sequence space to isolate a motifs’ context-free effect. By including the shuffled sequence, we aim to refine this by integrating over the motifs’ effect across compositions, effectively highlighting the component of importance due to ordered background context (hence, attempting to re-tain information from surrounding, non-shuffled motifs). Nonetheless, when searching for epistatic effects, the same issue as the original global importance persists—by taking the average across all sequences the metric still effectively integrates out sequence-specific effects. Accordingly, we posit that averaging over the sequence space is at odds with learning sequence-specific (i.e., epistatic) effects, and therefore suggest using the variance in local importance as a proxy for motif context-specificity:

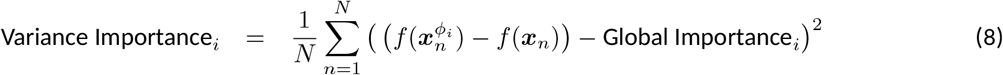

Intuitively, when a motif has context-specific effects in a given sequence, the distance between the sequence’s local importance will differ from the global, population-averaged importance. These distances can therefore help highlight interactions between an implanted motif and the sequence to which it was injected, either due to interactions with other motifs or other, spurious sequence elements. Explicitly, apart from other motifs influencing the implanted motif’s effect, this metric can be confounded by sequence elements (e.g. local GC content), for which we also examined using as a second-moment estimator the variance in local importance across the shuffled sequences, as well as the variance of the delta importance in attempt to remove the non-motif signal. Lastly, we considered whether the difference between the importance variance on shuffled and the importance variance on the observed sequences could elucidate any epistatic information. We examined first the implied covariance between the two quantities, hypothesizing that the importance variance in the observed sequences should be independent of the importance variance in the shuffled sequences (and therefore, equivalent) unless there is a non-sequence, non-compositional property of the motif (e.g., position in the sequence) affecting its importance:

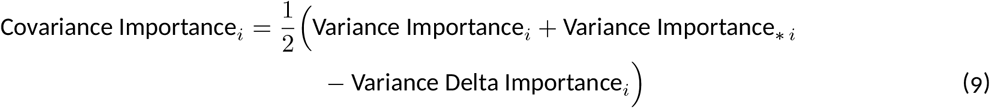

where Variance Importance_*i*_ is calculated as in equation 8 on the observed sequences, Variance Importance_∗ *i*_ is calculated as in equation 8 but using the shuffled sequences, and Variance Delta Importance_*i*_ is calculated over the variance in local delta importances shown in equation 7. Finally, to account for any first-moment leakage in our second-moment metrics, we also scored the Global Importance-residualized Variance Importance by regressing the Variance Importance (dependent variable) onto the Global Importance (independent variable) in a simple linear regression, and using the residuals.

For purposes of calculating AUROC, we used the absolute value of the GIA metrics so that any values away from 0 were predictive of motifs involved with epistasis. For calculating top-1 precision in the half-additive and half-epistatic context, we examined only discovered motifs (which is the set of motifs for which we ran GIA) and calculated across simulation realizations for a given parameter setting, architecture, and xAI model how frequently the top-GIA implied motif was involved in an epistatic interaction (precision), the proportion of discovered motifs that were involved in an epistatic interaction (prevalence), and the enrichment as the mean ratio of the precision over the prevalence. For the experimental data, we ran a regression of the discovered motifs’ second-moment importance onto the absolute first-moment importance, and prioritized motifs based on the value of their residuals (where, positive implied greater enrichment of second-moment importance relative to first-moment).

## References

[1] Rajiv Movva, Peyton Greenside, Georgi K. Marinov, Surag Nair, Avanti Shrikumar, and Anshul Kundaje. Deciphering regulatory dna sequences and noncoding genetic variants using neural network models of massively parallel reporter assays. PLOS ONE, 14(6):e0218073, 2019.

[2] Justin B. Kinney and David M. McCandlish. Massively parallel assays and quantitative sequence–function relation-ships. Annual Review of Genomics and Human Genetics, 20(1):99–127, August 2019.

[3] Shengdong Ke, Vincent Anquetil, Jorge Rojas Zamalloa, Alisha Maity, Anthony Yang, Mauricio A. Arias, Sergey Kalachikov, James J. Russo, Jingyue Ju, and Lawrence A. Chasin. Saturation mutagenesis reveals manifold determinants of exon definition. Genome Research, 28(1):11–24, December 2017.

[4] Christian B. Macdonald, David Nedrud, Patrick Rockefeller Grimes. Donovan Trinidad, James S. Fraser, and Willow Coyote-Maestas. Dimple: deep insertion, deletion, and missense mutation libraries for exploring protein variation in evolution, disease, and biology. Genome Biology, 24(1), February 2023.

[5] Aaron N. Brooks, Amanda L. Hughes, Sandra Clauder-Münster, Leslie A. Mitchell, Jef D. Boeke, and Lars M. Steinmetz. Transcriptional neighborhoods regulate transcript isoform lengths and expression levels. Science, 375(6584):1000–1005, March 2022.

[6] Jonas Koeppel, Raphael Ferreira, Thomas Vanderstichele, Lisa Maria Riedmayr, Elin Madli Peets, Gareth Girling, Juliane Weller, Pierre Murat, Fabio Giuseppe Liberante, Tom Ellis, George McDonald Church, and Leopold Parts. Randomizing the human genome by engineering recombination between repeat elements. Science, 387(6733), January 2025.

[7] Susan E. Liao, Mukund Sudarshan, and Oded Regev. Deciphering rna splicing logic with interpretable machine learning. Proceedings of the National Academy of Sciences, 120(41):e2221165120, 2023.

[8] Sebastian M. Castillo-Hair and Georg Seelig. Machine learning for designing next-generation mrna therapeutics. Accounts of Chemical Research, 55(1):24–34, 2022. PMID: 34905691.

[9] Abdul Muntakim Rafi, Daria Nogina, Dmitry Penzar, Dohoon Lee, Danyeong Lee, Nayeon Kim, Sangyeup Kim, Dohyeon Kim, Yeojin Shin, Il-Youp Kwak, Georgy Meshcheryakov, Andrey Lando, Arsenii Zinkevich, ByeongChan Kim, Juhyun Lee, Taein Kang, Eeshit Dhaval Vaishnav, Payman Yadollahpour, Random Promoter DREAM Challenge Consortium, Sun Kim, Jake Albrecht, Aviv Regev, Wuming Gong, Ivan V. Kulakovskiy, Pablo Meyer, and Carl de Boer. Evaluation and optimization of sequence-based gene regulatory deep learning models. April 2023.

[10] Alexander B. Rosenberg, Rupali P. Patwardhan, Jay Shendure, and Georg Seelig. Learning the sequence determinants of alternative splicing from millions of random sequences. Cell, 163(3):698–711, October 2015.

[11] Jian Zhou and Olga G Troyanskaya. Predicting effects of noncoding variants with deep learning-based sequence model. Nat. Methods, 12(10):931–934, October 2015.

[12] Babak Alipanahi, Andrew Delong, Matthew T Weirauch, and Brendan J Frey. Predicting the sequence specificities of DNA- and RNA-binding proteins by deep learning. Nat. Biotechnol., 33(8):831–838, August 2015.

[13] David R. Kelley, Jasper Snoek, and John L. Rinn. Basset: learning the regulatory code of the accessible genome with deep convolutional neural networks. Genome Research, 26(7):990–999, 2016.

[14] Gherman Novakovsky, Nick Dexter, Maxwell W. Libbrecht, Wyeth W. Wasserman, and Sara Mostafavi. Obtaining genetics insights from deep learning via explainable artificial intelligence. Nature Reviews Genetics, 24(2):125–137, October 2022.

[15] Masayuki Nagai, Alan E. Murphy, Kaeli Rizzo, and Peter K. Koo. Toward interpretable and generalizable ai in regulatory genomics, 2026.

[16] Peter K. Koo and Matt Ploenzke. Deep learning for inferring transcription factor binding sites. Current Opinion in Systems Biology, 19:16–23, 2020.

[17] Julia Zeitlinger, Sushmita Roy, Ferhat Ay, Anthony Mathelier, Alejandra Medina-Rivera, Shaun Mahony, Saurabh Sinha, and Jason Ernst. Perspective on recent developments and challenges in regulatory and systems genomics. Bioinformatics Advances, 5(1), December 2024.

[18] Valerie Chen, Muyu Yang, Wenbo Cui, Joon Sik Kim, Ameet Talwalkar, and Jian Ma. Applying interpretable machine learning in computational biology—pitfalls, recommendations and opportunities for new developments. Nature Methods, 21(8):1454–1461, August 2024.

[19] Peter K. Koo and Sean R. Eddy. Representation learning of genomic sequence motifs with convolutional neural networks. PLOS Computational Biology, 15(12):1–17, 12 2019.

[20] Gonzalo Benegas, Sanjit Singh Batra, and Yun S. Song. Dna language models are powerful predictors of genomewide variant effects. Proceedings of the National Academy of Sciences, 120(44), October 2023.

[21] Yanrong Ji, Zhihan Zhou, Han Liu, and Ramana V Davuluri. Dnabert: pre-trained bidirectional encoder representations from transformers model for dna-language in genome. Bioinformatics, 37(15):2112–2120, February 2021.

[22] Pooja Kathail, Richard W. Shuai, Ryan Chung, Chun Jimmie Ye, Gabriel B. Loeb, and Nilah M. Ioannidis. Current genomic deep learning models display decreased performance in cell type-specific accessible regions. Genome Biology, 25(1), August 2024.

[23] Kishore Jaganathan, Sofia Kyriazopoulou Panagiotopoulou, Jeremy F. McRae, Siavash Fazel Darbandi, David Knowles, Yang I. Li, Jack A. Kosmicki, Juan Arbelaez, Wenwu Cui, Grace B. Schwartz, Eric D. Chow, Efstathios Kanterakis, Hong Gao, Amirali Kia, Serafim Batzoglou, Stephan J. Sanders, and Kyle Kai-How Farh. Predicting splicing from primary sequence with deep learning. Cell, 176(3):535–548.e24, January 2019.

[24] Ziga Avsec, Vikram Agarwal, Daniel Visentin, Joseph R. Ledsam, Agnieszka Grabska-Barwinska, Kyle R. Taylor, Yannis Assael, John Jumper, Pushmeet Kohli, and David R. Kelley. Effective gene expression prediction from sequence by integrating long-range interactions. Nature Methods, 18(10):1196–1203, October 2021.

[25] Ziga Avsec, Natasha Latysheva, Jun Cheng, Guido Novati, Kyle R. Taylor, Tom Ward, Clare Bycroft, Lauren Nicolaisen, Eirini Arvaniti, Joshua Pan, Raina Thomas, Vincent Dutordoir, Matteo Perino, Soham De, Alexander Karollus, Adam Gayoso, Toby Sargeant, Anne Mottram, Lai Hong Wong, Pavol Drotár, Adam Kosiorek, Andrew Senior, Richard Tanburn, Taylor Applebaum, Souradeep Basu, Demis Hassabis, and Pushmeet Kohli. Advancing regulatory variant effect prediction with alphagenome. Nature, 649(8099):1206–1218, January 2026.

[26] Scott Lundberg and Su-In Lee. A unified approach to interpreting model predictions, 2017.

[27] Avanti Shrikumar, Peyton Greenside, and Anshul Kundaje. Learning important features through propagating activation differences. 2017.

[28] Avanti Shrikumar, Katherine Tian, Ziga Avsec, Anna Shcherbina, Abhimanyu Banerjee, Mahfuza Sharmin, Surag Nair, and Anshul Kundaje. Technical note on transcription factor motif discovery from importance scores (tf-modisco) version 0.5.6.5, 2018.

[29] Eva I. Prakash, Avanti Shrikumar, and Anshul Kundaje. Towards more realistic simulated datasets for benchmarking deep learning models in regulatory genomics. In David A. Knowles, Sara Mostafavi, and Su-In Lee, editors, Proceedings of the 16th Machine Learning in Computational Biology meeting, volume 165 of Proceedings of Machine Learning Research, pages 58–77. PMLR, 22–23 Nov 2022.

[30] Ling Chen and John A. Capra. Learning and interpreting the gene regulatory grammar in a deep learning framework. PLOS Computational Biology, 16(11):e1008334, November 2020.

[31] Katalin Ferenc, Lorenzo Martini, Ieva Rauluseviciute, Geir Kjetil Ferkingstad Sandve, and Anthony Mathelier. inmotifin: a lightweight end-to-end simulation software for regulatory sequences. Bioinformatics, 42(2), January 2026.

[32] Antonio Majdandzic, Chandana Rajesh, and Peter K. Koo. Correcting gradient-based interpretations of deep neural networks for genomics. Genome Biology, 24(1), May 2023.

[33] Fahad Ullah and Asa Ben-Hur. A self-attention model for inferring cooperativity between regulatory features. Nucleic Acids Research, 49(13):e77.–e77, 05 2021.

[34] Jiawei Li, Yuqian Pu, Jijun Tang, Quan Zou, and Fei Guo. Deepatt: a hybrid category attention neural network for identifying functional effects of dna sequences. Briefings in Bioinformatics, 22(3), August 2020.

[35] Tony Zeng and Yang I Li. Predicting rna splicing from dna sequence using pangolin. Genome Biology, 23(1), April 2022.

[36] Xin Cheng, Jun Wang, Qianyue Li, and Taigang Liu. Bilstm-5mc: A bidirectional long short-term memory-based approach for predicting 5-methylcytosine sites in genome-wide dna promoters. Molecules, 26(24):7414, December 2021.

[37] Peter K. Koo and Matt Ploenzke. Improving representations of genomic sequence motifs in convolutional networks with exponential activations. Nature Machine Intelligence, 3(3):258–266, February 2021.

[38] Kevin B. Dsouza, Alexandra Maslova, Ediem Al-Jibury, Matthias Merkenschlager, Vijay K. Bhargava, and Maxwell W. Libbrecht. Learning representations of chromatin contacts using a recurrent neural network identifies genomic drivers of conformation. Nature Communications, 13(1), June 2022.

[39] Kaiming He, Xiangyu Zhang, Shaoqing Ren, and Jian Sun. Deep residual learning for image recognition. In 2016 IEEE Conference on Computer Vision and Pattern Recognition (CVPR), pages 770–778, 2016.

[40] Robert Frömel, Julia Rühle, Aina Bernal Martinez, Chelsea Szu-Tu, Felix Pacheco Pastor, Rosa Martinez-Corral, and Lars Velten. Design principles of cell-state-specific enhancers in hematopoiesis. Cell, 188(12):3202–3218.e21, June 2025. Publisher: Elsevier.

[41] Vikram Agarwal, Fumitaka Inoue, Max Schubach, Dmitry Penzar, Beth K. Martin, Pyaree Mohan Dash, Pia Keukeleire, Zicong Zhang, Ajuni Sohota, Jingjing Zhao, Ilias Georgakopoulos-Soares, William S. Noble, Galip Gürkan Yardımcı, Ivan V. Kulakovskiy, Martin Kircher, Jay Shendure, and Nadav Ahituv. Massively parallel characterization of transcriptional regulatory elements. Nature, 639(8054):411–420, January 2025.

[42] Eva I. Prakash, Avanti Shrikumar, and Anshul Kundaje. Towards more realistic simulated datasets for benchmarking deep learning models in regulatory genomics. In David A. Knowles, Sara Mostafavi, and Su-In Lee, editors, Proceedings of the 16th Machine Learning in Computational Biology meeting, volume 165 of Proceedings of Machine Learning Research, pages 58–77. PMLR, 22–23 Nov 2022.

[43] Peter K. Koo, Antonio Majdandzic, Matthew Ploenzke, Praveen Anand, and Steffan B. Paul. Global importance analysis: An interpretability method to quantify importance of genomic features in deep neural networks. PLOS Computational Biology, 17(5):e1008925, May 2021.

[44] Mike Thompson, Mariano Martín, Trinidad Sanmartín Olmo, Chandana Rajesh, Peter K. Koo, Benedetta Bolognesi, and Ben Lehner. Massive experimental quantification allows interpretable deep learning of protein aggregation. Science Advances, 11(18):eadt5111, 2025.

[45] Ryan Z. Friedman, Avinash Ramu, Sara Lichtarge, Yawei Wu, Lloyd Tripp, Daniel Lyon, Connie A. Myers, David M. Granas, Maria Gause, Joseph C. Corbo, Barak A. Cohen, and Michael A. White. Active learning of enhancers and silencers in the developing neural retina. Cell Systems, 16(1):101163, January 2025.

[46] Adam Klie, David Laub, James V. Talwar, Hayden Stites, Tobias Jores, Joe J. Solvason, Emma K. Farley, and Hannah Carter. Predictive analyses of regulatory sequences with eugene. Nature Computational Science, 3(11):946–956, November 2023.

[47] Bernardo P. de Almeida, Franziska Reiter, Michaela Pagani, and Alexander Stark. Deepstarr predicts enhancer activity from dna sequence and enables the de novo design of synthetic enhancers. Nature Genetics, 54(5):613–624, May 2022.

[48] Peyton Greenside, Tyler Shimko, Polly Fordyce, and Anshul Kundaje. Discovering epistatic feature interactions from neural network models of regulatory dna sequences. Bioinformatics, 34(17):i629–i637, 2018.

[49] Shushan Toneyan and Peter K. Koo. Interpreting cis-regulatory interactions from large-scale deep neural networks. Nature Genetics, 56(11):2517–2527, 2024.

[50] Timothy Fuqua and Andreas Wagner. De-novo promoters emerge more readily from random dna than from genomic dna. August 2025.

[51] Carl G de Boer and Jussi Taipale. Hold out the genome: a roadmap to solving the cis-regulatory code. Nature, 625(7993):41–50, January 2024.

[52] Oriol Fornes, Jaime A Castro-Mondragon, Aziz Khan, Robin van der Lee, Xi Zhang, Phillip A Richmond, Bhavi P Modi, Solenne Correard, Marius Gheorghe, Damir Baranasic, Walter Santana-Garcia, Ge Tan, Jeanne Chèneby, Benoit Ballester, François Parcy, Albin Sandelin, Boris Lenhard, Wyeth W Wasserman, and Anthony Mathelier. Jaspar 2020: update of the open-access database of transcription factor binding profiles. Nucleic Acids Research, 48(D1):D87–D92, 11 2019.

[53] Peter M. Visscher, William G. Hill, and Naomi R. Wray. Heritability in the genomics era — concepts and misconceptions. Nature Reviews Genetics, 9(4):255–266, March 2008.

[54] Vikram Agarwal, Fumitaka Inoue, Max Schubach, Beth K. Martin, Pyaree Mohan Dash, Zicong Zhang, Ajuni Sohota, William Stafford Noble, Galip Gürkan Yardimci, Martin Kircher, Jay Shendure, and Nadav Ahituv. Massively parallel characterization of transcriptional regulatory elements in three diverse human cell types. March 2023.

[55] Shobhit Gupta, John A Stamatoyannopoulos, Timothy L Bailey, and William Stafford Noble. Quantifying similarity between motifs. Genome Biology, 8(2), February 2007.

