## Supplementary Figures for "Learning causal regulatory motifs and grammars using deep learning models and massively parallel reporter assays"

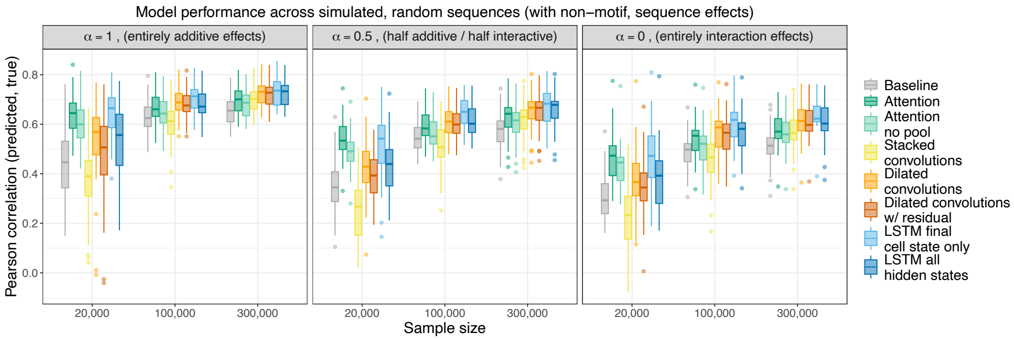


Figure S1: **Model performance in simulated data where non-motif effects contribute to phenotypic variability.** In addition to varying motif additivity, these simulation runs include a non-motif effect (Methods) that accounts for 33% of phenotypic variability. As LSTMs can more readily learn this effect, the gap between them and attention-based models at low samples is less dramatic than in the absence of such an effect (where attention is the most efficient at learning motif-based grammars).

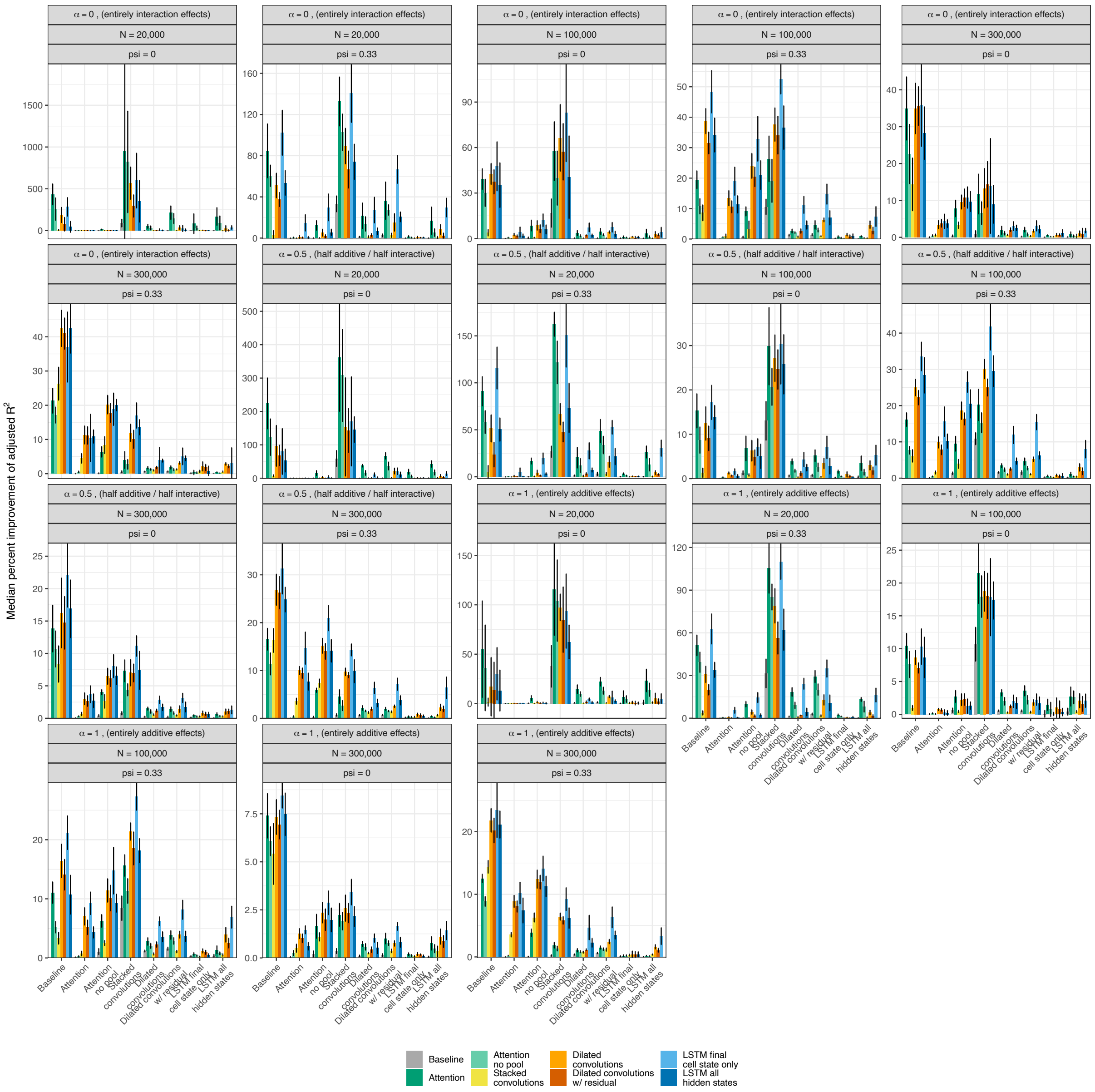


Figure S2: **Increase in variability explained by different models over the test set.** To evaluate whether models captured independent phenotypic variability in our simulations, we used a linear model between the true phenotype and neural network predictions to calculate the adjusted $R^2$ per architecture (x-axis). We then added to the linear model the prediction of one additional model (color) in a round robin fashion. In other words, we calculated a base adjusted $R^2$ for one model, then included at most in the linear model two predictors—one from the base model, and another from the tested (and plotted) model. This enables us to evaluate whether similarly performing models capture different components of the phenotypic variability. We plot here the median percent improvement (and 95% confidence intervals) for all neural networks that converged and explained a baseline adjusted $R^2$ of 0.05.

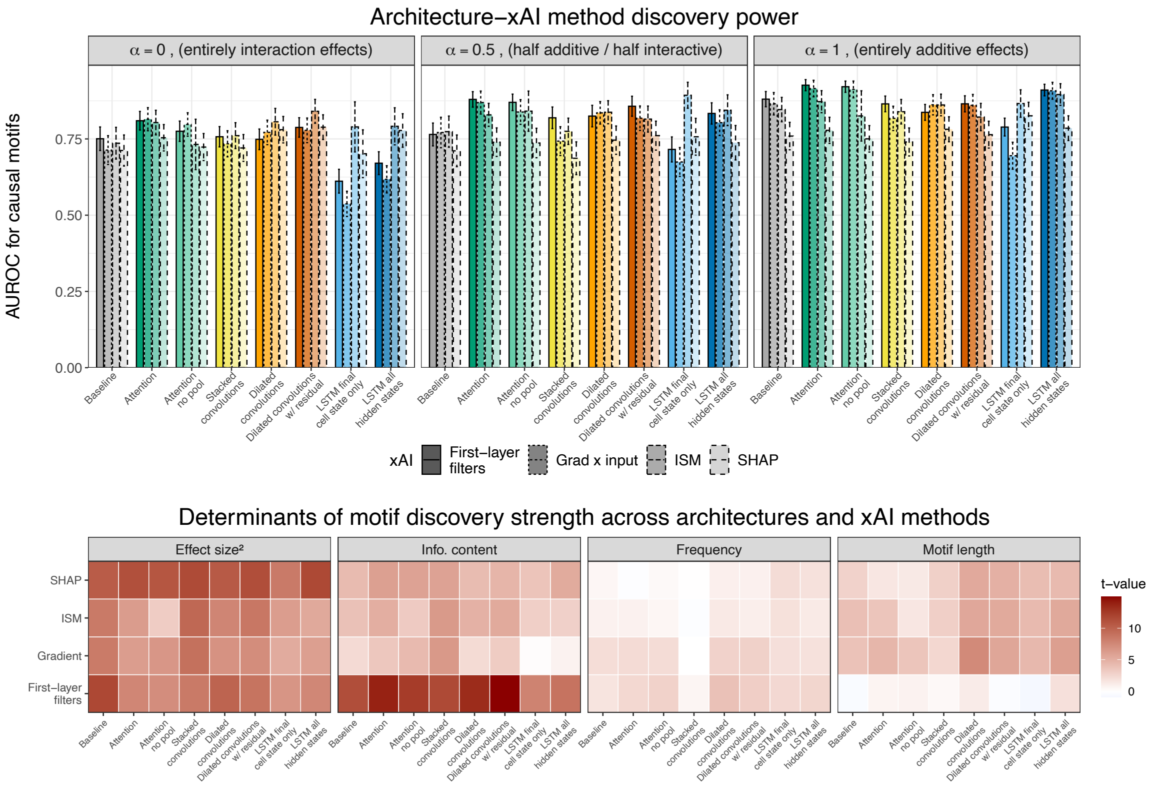


Figure S3: **Motif discovery characteristics with 300,000 sequences and no non-motif effects.** (A) The AUC of each model-xAI pair for prioritizing causal motifs across genetic architectures. Error bars represent 95% confidence intervals obtained via non-parametric bootstrapping. (B) Association statistics between motif characteristics and discovery strength for correctly identified causal motifs.

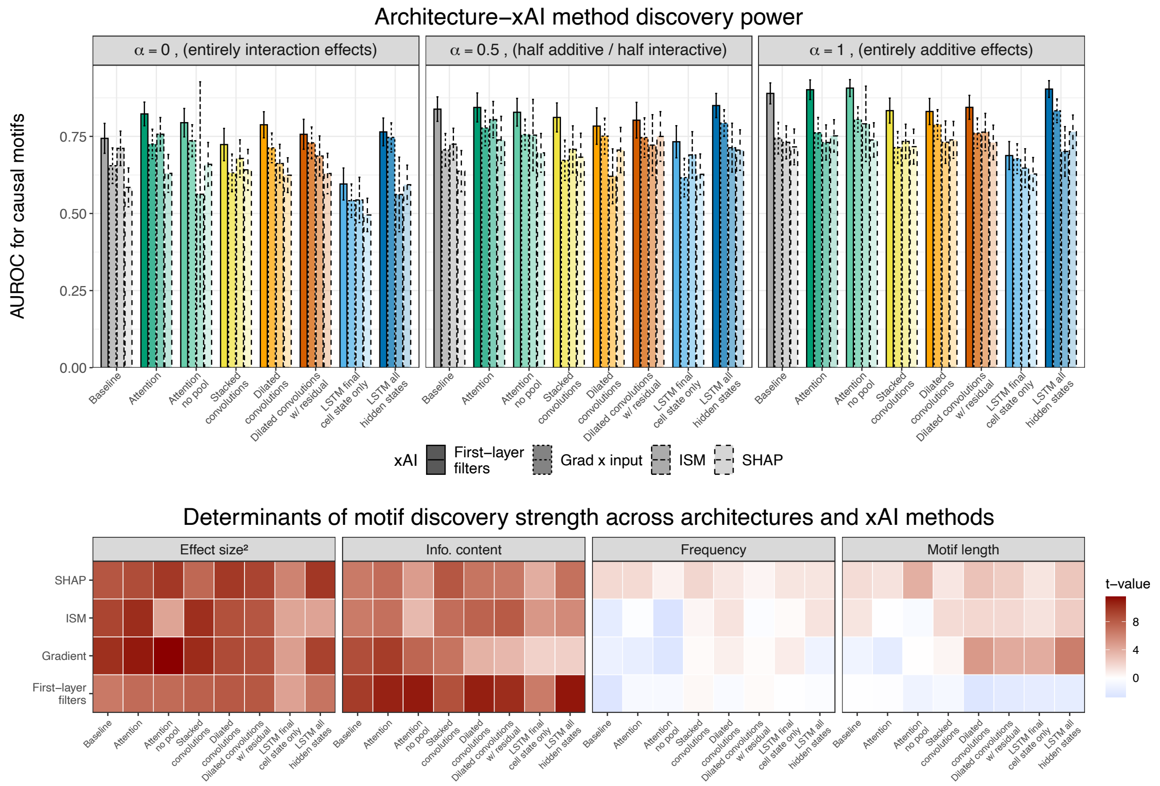


Figure S4: **Motif discovery characteristics with 300,000 sequences with non-motif effects.** (A) The AUC of each model-xAI pair for prioritizing causal motifs across genetic architectures. Error bars represent 95% confidence intervals obtained via non-parametric bootstrapping. (B) Association statistics between motif characteristics and discovery strength for correctly identified causal motifs.

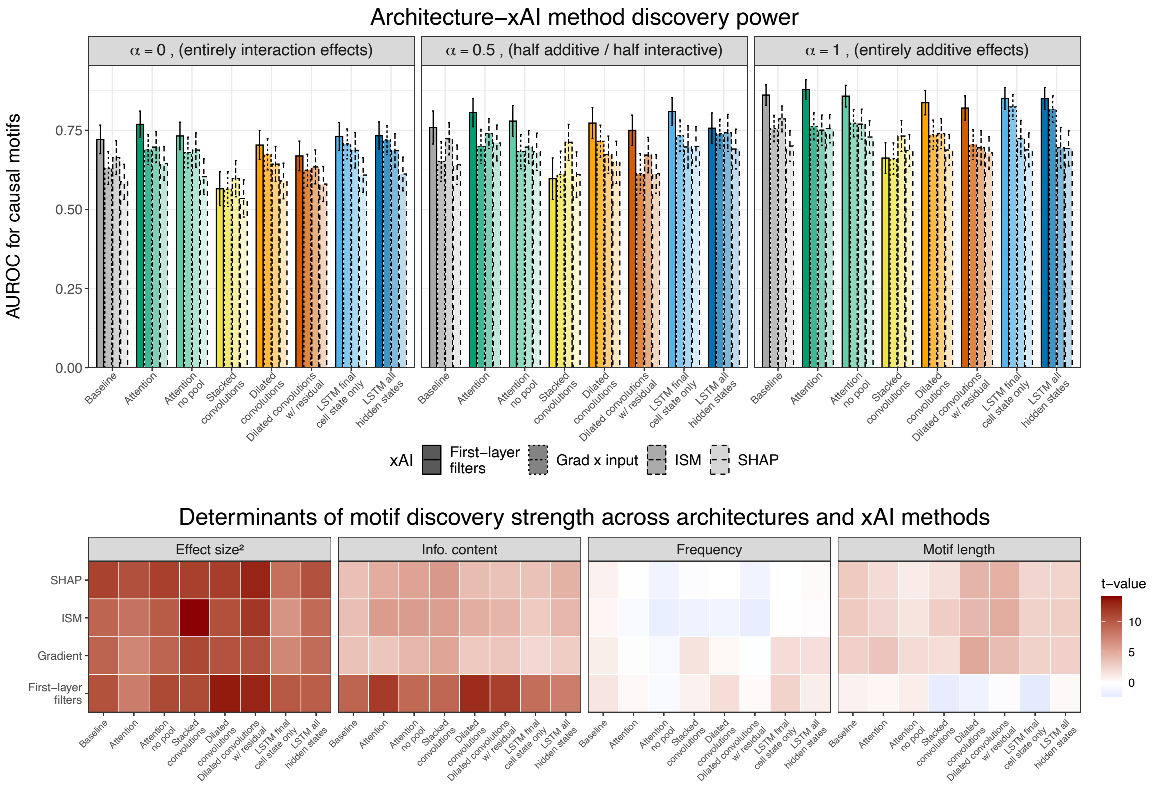


Figure S5: **Motif discovery characteristics with 100,000 sequences and no non-motif effects.** (A) The AUC of each model-xAI pair for prioritizing causal motifs across genetic architectures. Error bars represent 95% confidence intervals obtained via non-parametric bootstrapping. (B) Association statistics between motif characteristics and discovery strength for correctly identified causal motifs.

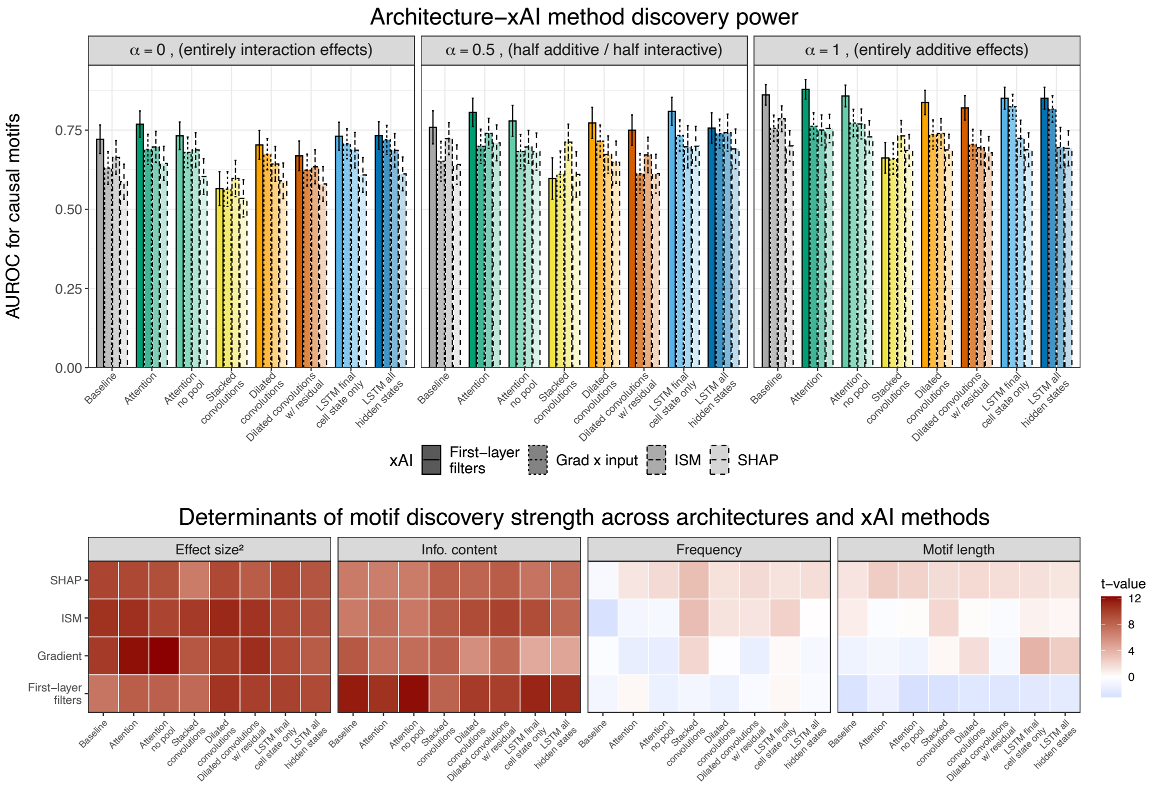


Figure S6: **Motif discovery characteristics with 100,000 sequences with non-motif effects.** (A) The AUC of each model-xAI pair for prioritizing causal motifs across genetic architectures. Error bars represent 95% confidence intervals obtained via non-parametric bootstrapping. (B) Association statistics between motif characteristics and discovery strength for correctly identified causal motifs.

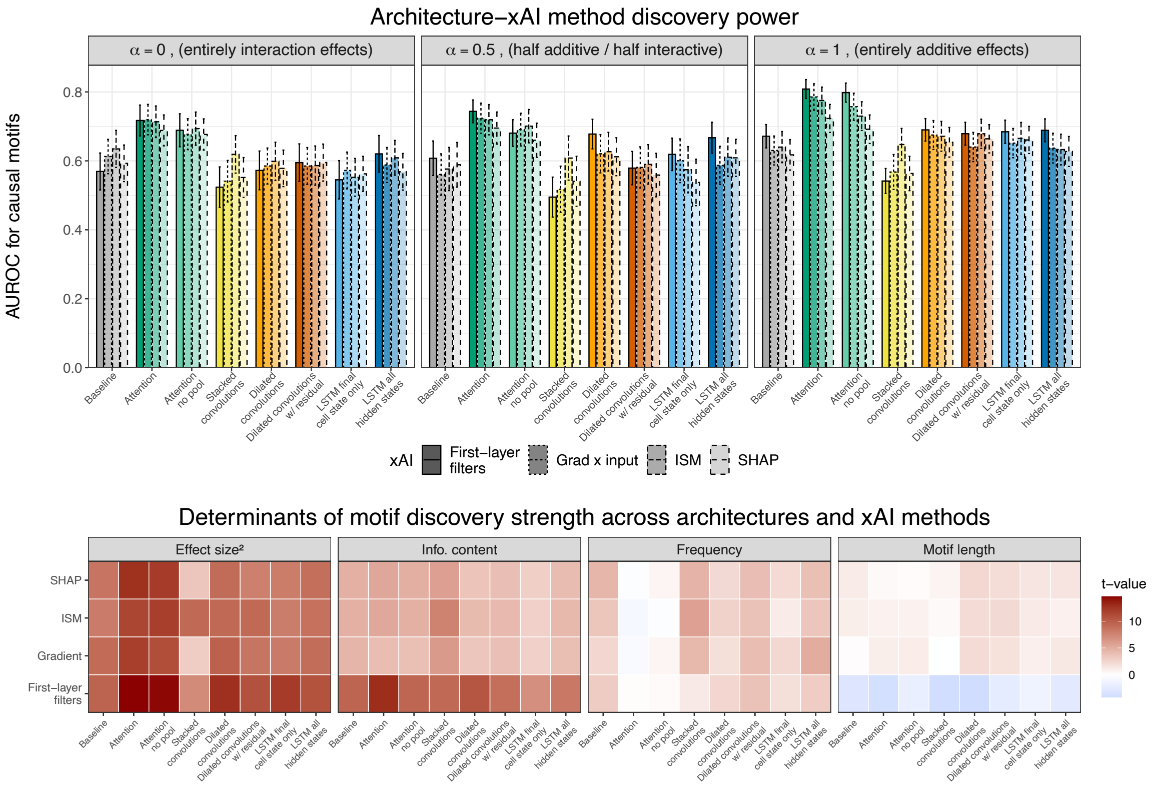


Figure S7: **Motif discovery characteristics with 20,000 sequences and no non-motif effects.** (A) The AUC of each model-xAI pair for prioritizing causal motifs across genetic architectures. Error bars represent 95% confidence intervals obtained via non-parametric bootstrapping. (B) Association statistics between motif characteristics and discovery strength for correctly identified causal motifs.

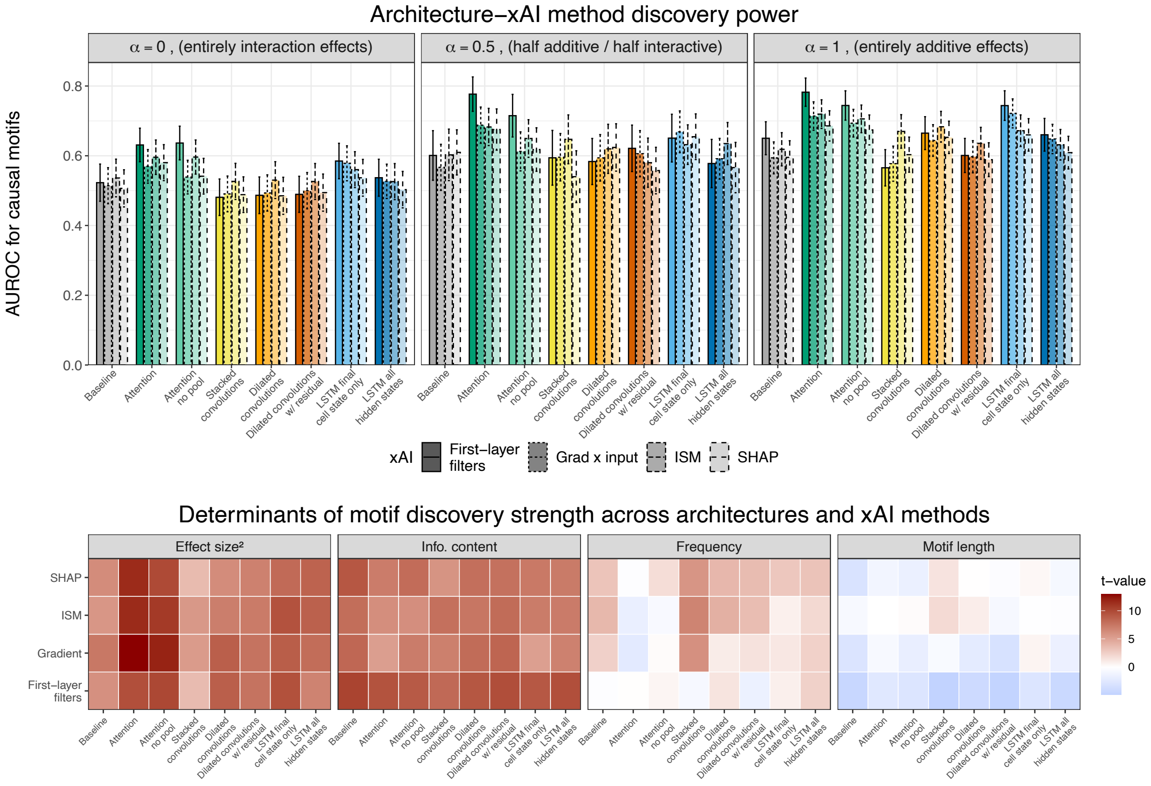


Figure S8: **Motif discovery characteristics with 20,000 sequences with non-motif effects.** (A) The AUC of each model-xAI pair for prioritizing causal motifs across genetic architectures. Error bars represent 95% confidence intervals obtained via non-parametric bootstrapping. (B) Association statistics between motif characteristics and discovery strength for correctly identified causal motifs.

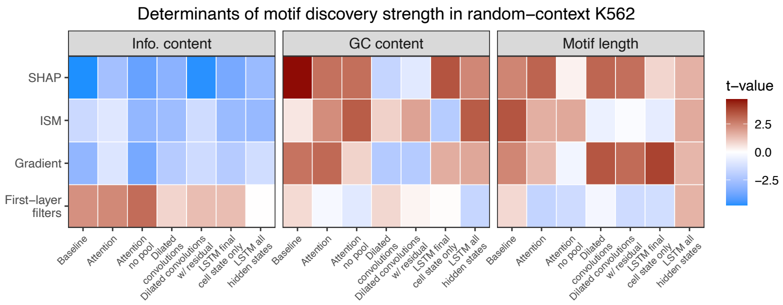


Figure S9: **Association of motif characteristics with discovery frequency in the Frömel et al. random sequences.** We ran a binomial regression of the number of times a motif was discovered given the number of times a model training instance converged, where motif information content, GC content, and length were included as explanatory variables. Regressions were run within a given architecture-xAI combination, and we included only motifs that were discovered at least across 2 model training instances to reduce spurious associations.

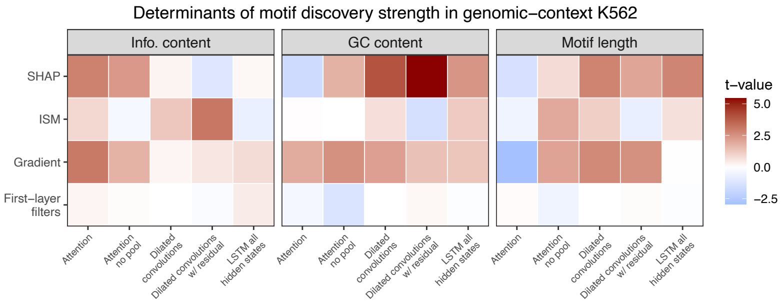


Figure S10: **Association of motif characteristics with discovery frequency in the Agarwal et al. genomic sequences in K562.** We ran a binomial regression of the number of times a motif was discovered given the number of times a model training instance converged, where motif information content, GC content, and length were included as explanatory variables. Regressions were run within a given architecture-xAI combination, and we included only motifs that were discovered at least across 2 model training instances to reduce spurious associations.

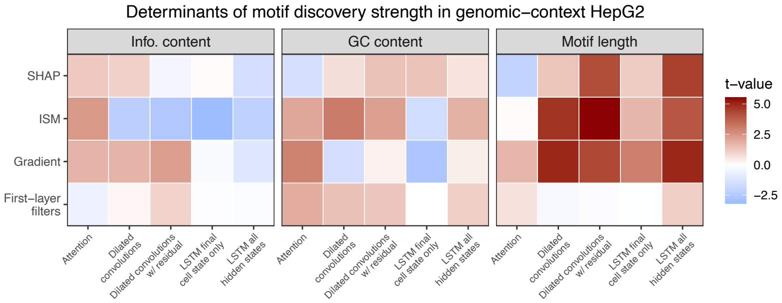


Figure S11: **Association of motif characteristics with discovery frequency in the Agarwal et al. genomic sequences in HepG2.** We ran a binomial regression of the number of times a motif was discovered given the number of times a model training instance converged, where motif information content, GC content, and length were included as explanatory variables. Regressions were run within a given architecture-xAI combination, and we included only motifs that were discovered at least across 2 model training instances to reduce spurious associations.

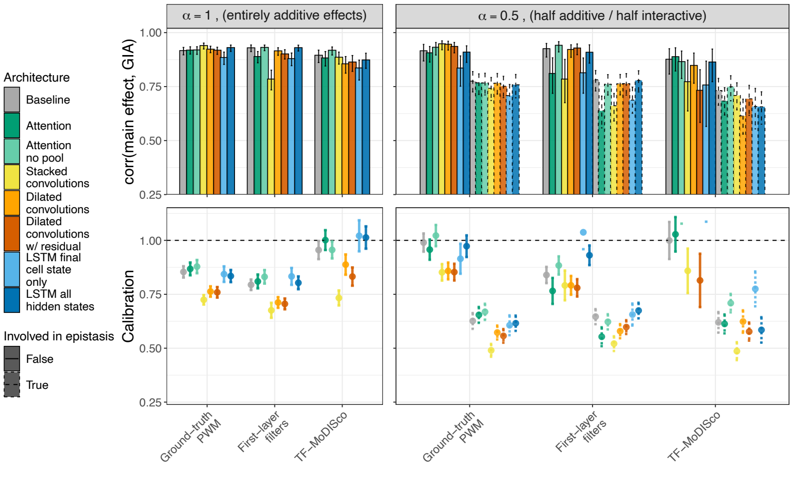


Figure S12: **Calibration of GIA to discover ground-truth main effects at N=300,000 and where motif effects explain all phenotypic variability.** Spearman correlation between true main effect and GIA-estimated effects are plotted in the upper panels, and calibration (slope from regression) of GIA estimation of the true effect is shown in the bottom.

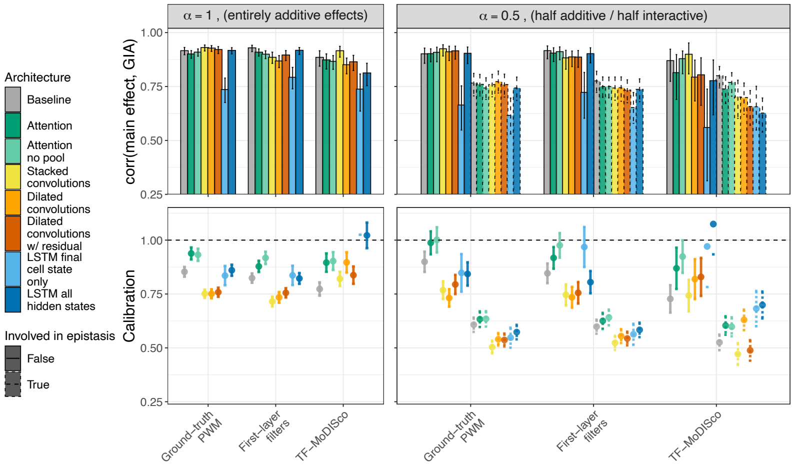


Figure S13: **Calibration of GIA to discover ground-truth main effects at N=300,000 and where non-motif effects explain 33% of phenotypic variability.** Spearman correlation between true main effect and GIA-estimated effects are plotted in the upper panels, and calibration (slope from regression) of GIA estimation of the true effect is shown in the bottom.

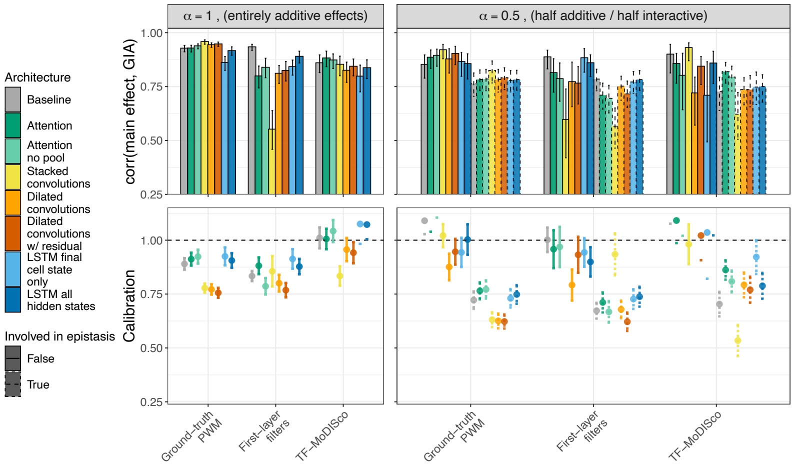


Figure S14: **Calibration of GIA to discover ground-truth main effects at N=100,000 and where motif effects explain all phenotypic variability.** Spearman correlation between true main effect and GIA-estimated effects are plotted in the upper panels, and calibration (slope from regression) of GIA estimation of the true effect is shown in the bottom.

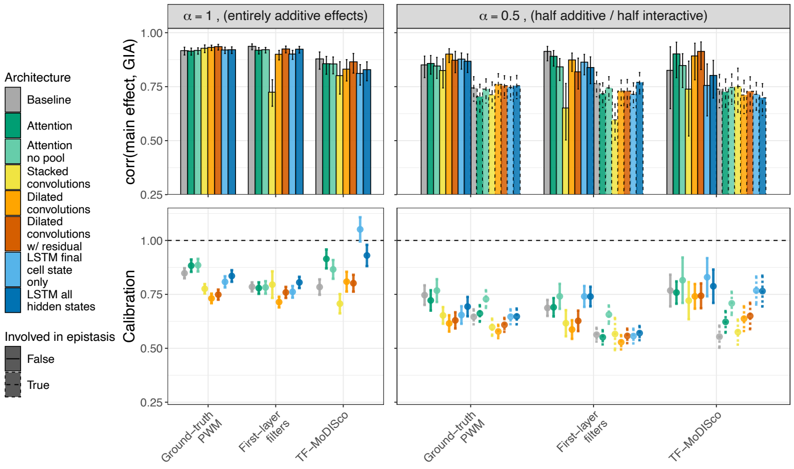


Figure S15: **Calibration of GIA to discover ground-truth main effects at N=100,000 and where non-motif effects explain 33% of phenotypic variability.** Spearman correlation between true main effect and GIA-estimated effects are plotted in the upper panels, and calibration (slope from regression) of GIA estimation of the true effect is shown in the bottom.

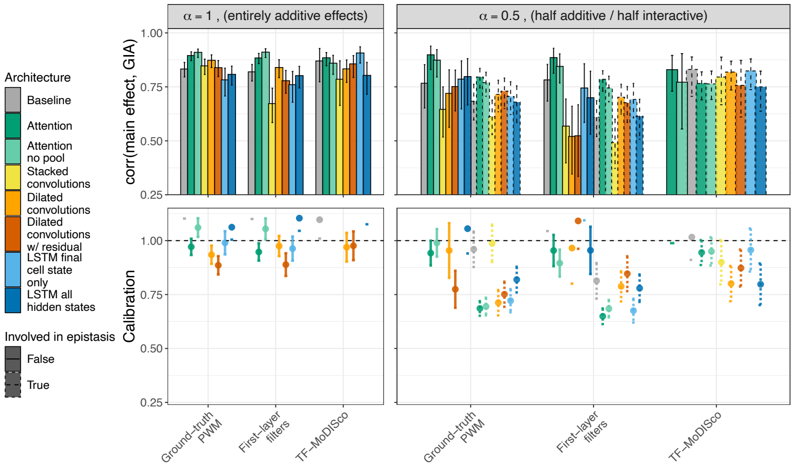


Figure S16: **Calibration of GIA to discover ground-truth main effects at N=20,000 and where motif effects explain all phenotypic variability.** Spearman correlation between true main effect and GIA-estimated effects are plotted in the upper panels, and calibration (slope from regression) of GIA estimation of the true effect is shown in the bottom. Missing method-xAI pairs represent combinations where fewer than 30 observations were discovered (the model was underpowered) and hence excluded from analysis.

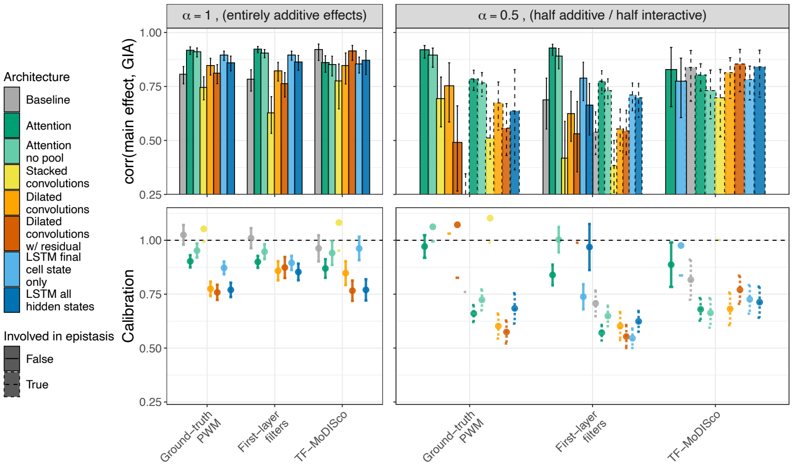


Figure S17: **Calibration of GIA to discover ground-truth main effects at N=20,000 and where non-motif effects explain 33% of phenotypic variability.** Spearman correlation between true main effect and GIA-estimated effects are plotted in the upper panels, and calibration (slope from regression) of GIA estimation of the true effect is shown in the bottom. Missing method-xAI pairs represent combinations where fewer than 30 observations were discovered (the model was underpowered) and hence excluded from analysis.

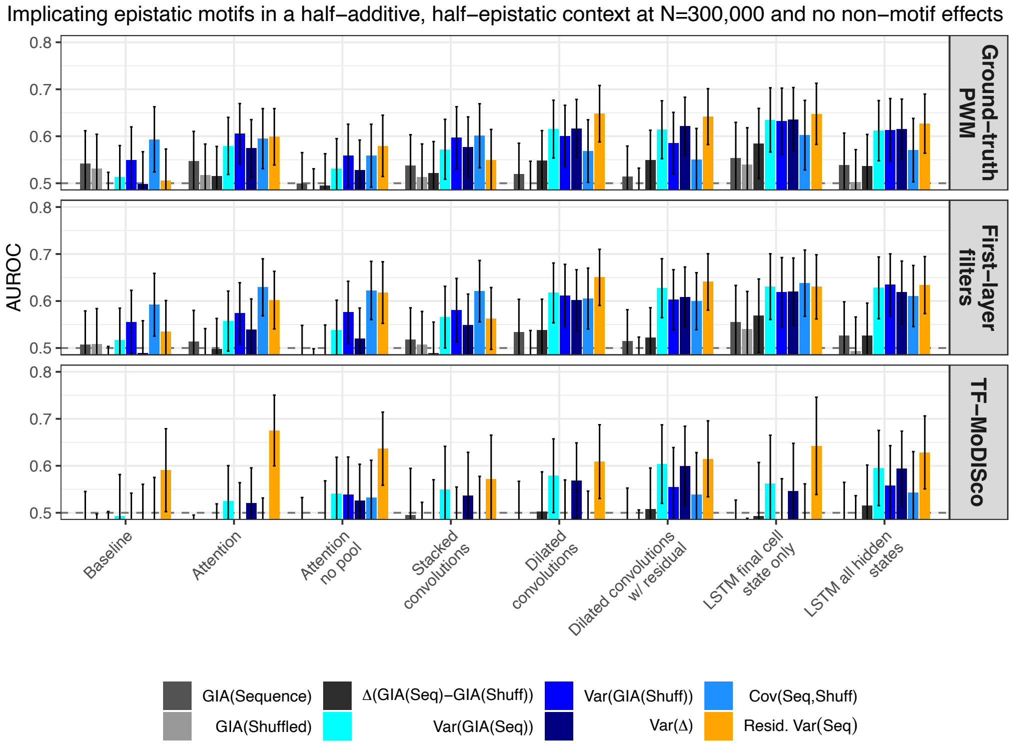


Figure S18: **Implicating epistatic motifs at N=300,000 and no non-motif sequence effects.** Here we examine the power of xAI-architecture combinations and various GIA metrics to classify whether a discovered motif is involved in an epistatic interaction or not. Any motif involved in any type of interaction (simple, upstream, distance or high-order) will be labeled 1, otherwise it will be labeled 0.

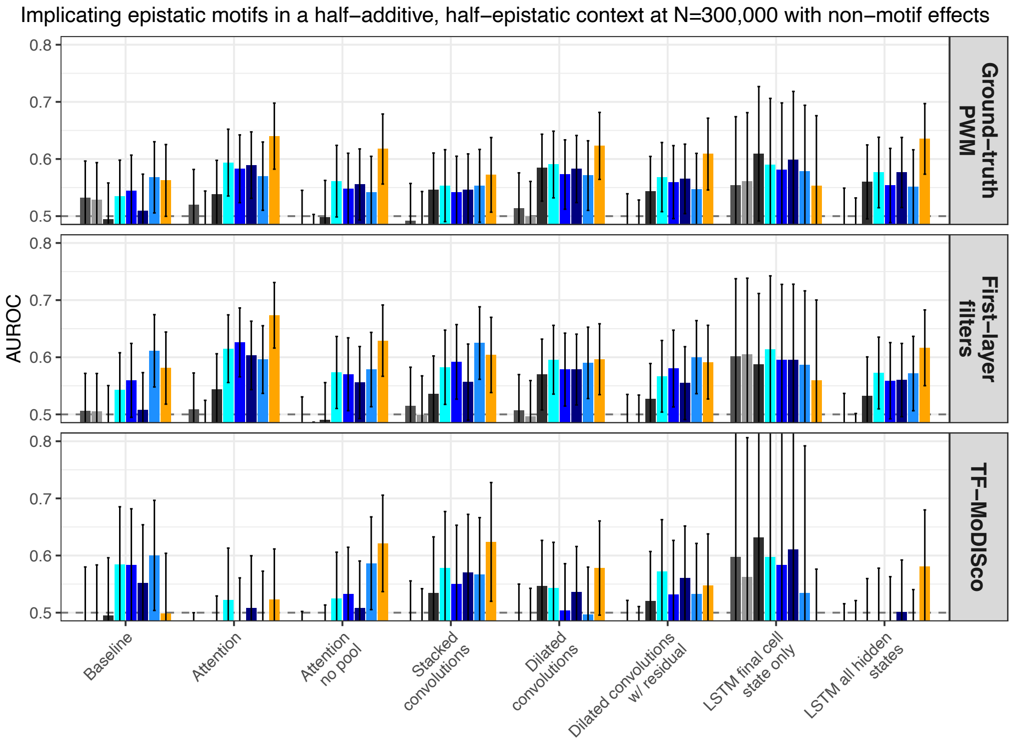


Figure S19: **Implicating epistatic motifs at N=300,000 in the presence of non-motif sequence effects.** Here we examine the power of xAI-architecture combinations and various GIA metrics to classify whether a discovered motif is involved in an epistatic interaction or not. Any motif involved in any type of interaction (simple, upstream, distance or high-order) will be labeled 1, otherwise it will be labeled 0.

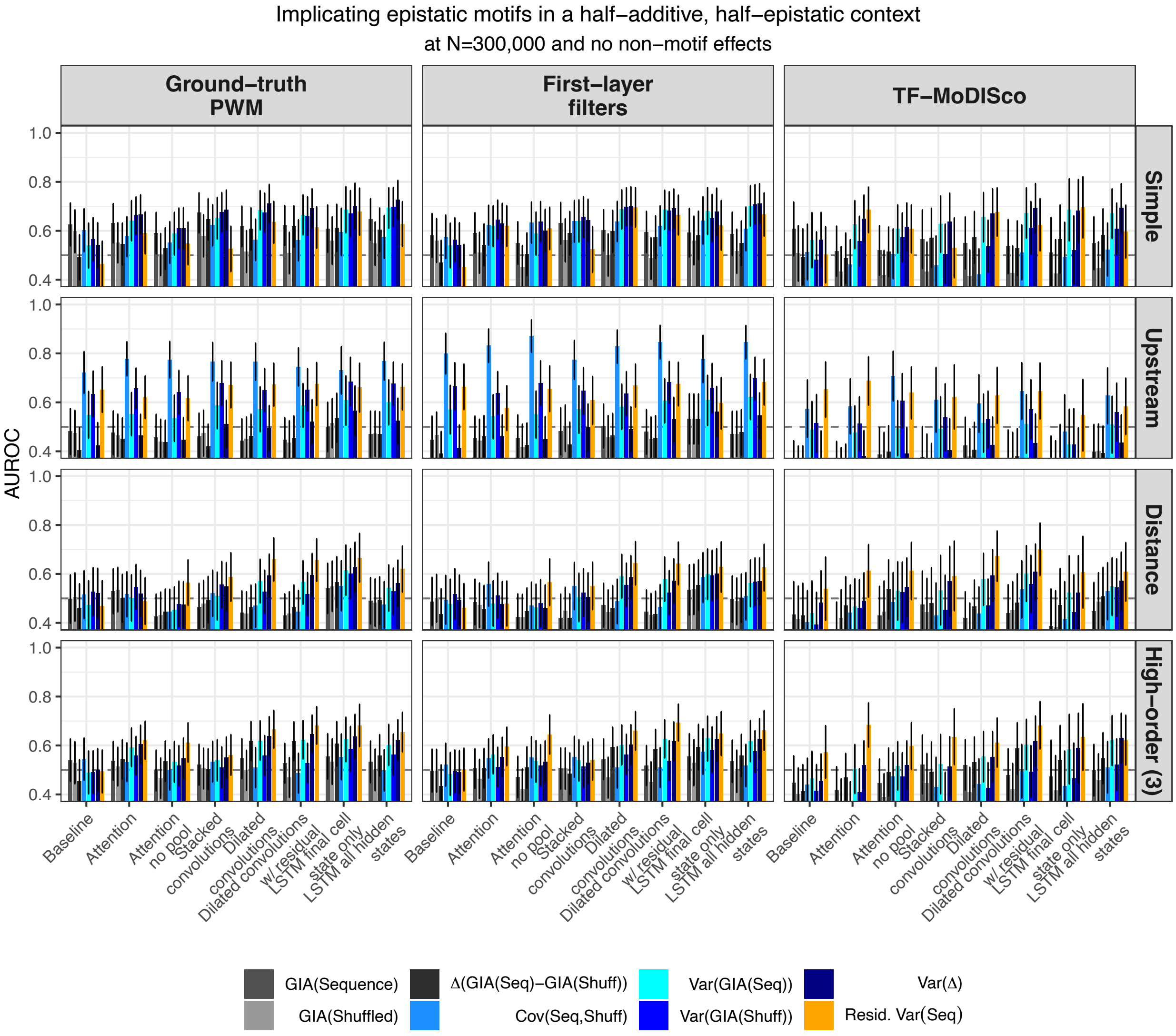


Figure S20: **Implicating epistatic motifs at N=300,000 in the absence of non-motif sequence effects.** Here we examine the power of xAI-architecture combinations and various GIA metrics to classify whether a discovered motif is involved in a specific epistatic interaction or not. For a given interaction class, any motif whose maximum interaction effect corresponding to that class (simple, upstream, distance or high-order) were labeled 1, and all other discovered motifs that were not involved with any interaction were labeled 0.

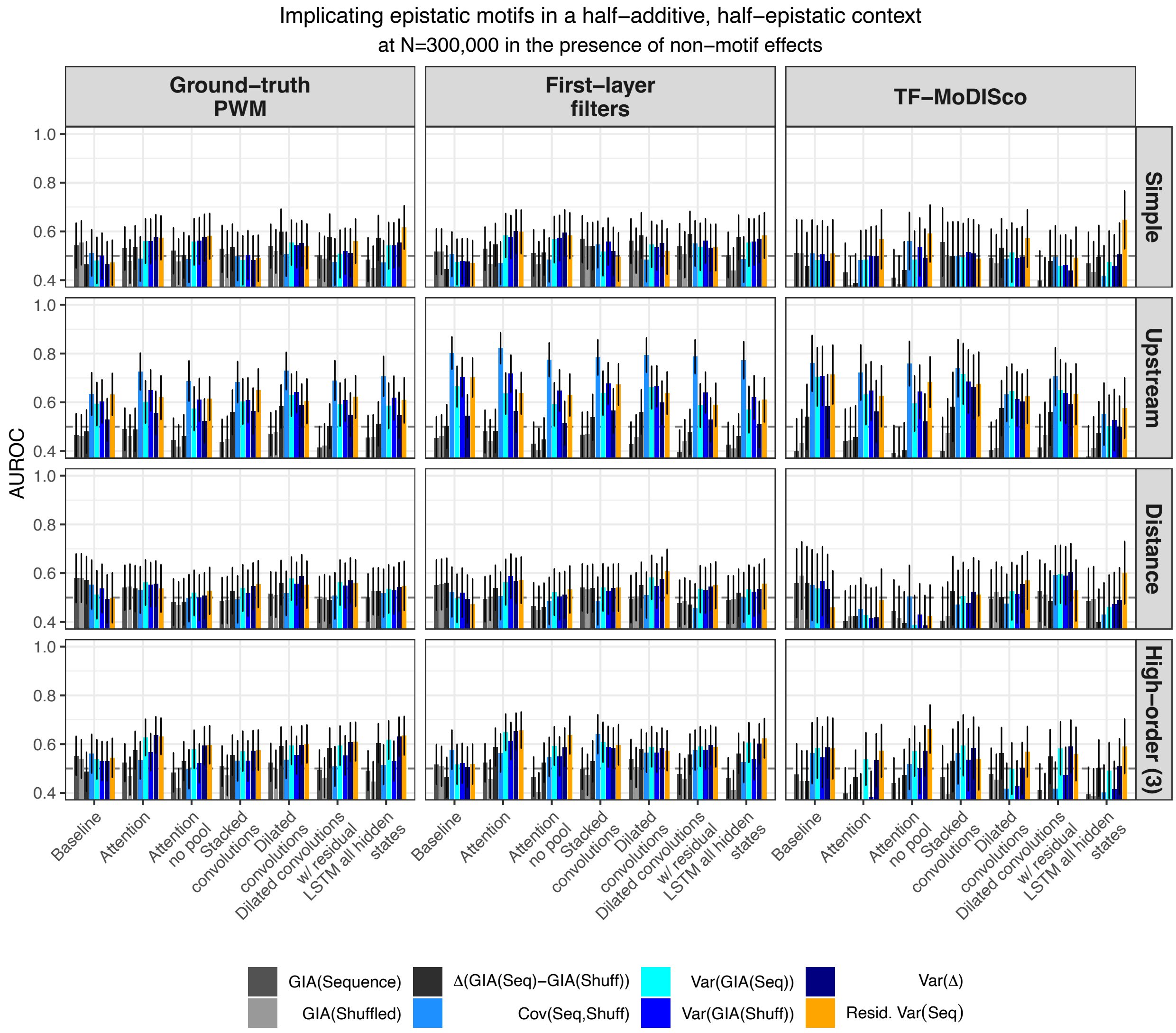


Figure S21: **Implicating epistatic motifs at N=300,000 in the presence of non-motif sequence effects.** Here we examine the power of xAI-architecture combinations and various GIA metrics to classify whether a discovered motif is involved in a specific epistatic interaction or not when 33% of the variability in phenotype due to sequence characteristics is made up of non-motif effects. For a given interaction class, any motif whose maximum interaction effect corresponding to that class (simple, upstream, distance or high-order) were labeled 1, and all other discovered motifs that were not involved with any interaction were labeled 0.

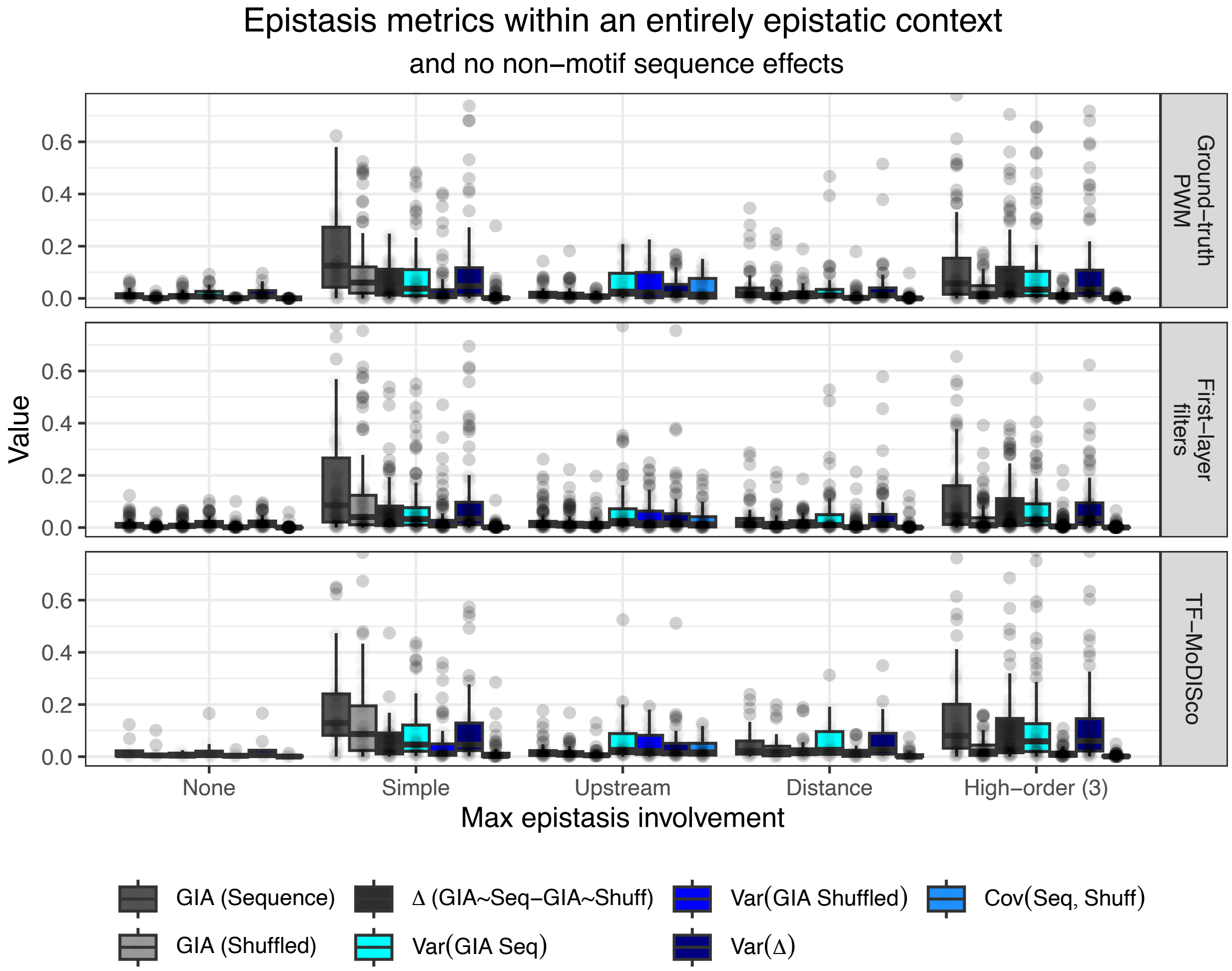


Figure S22: **GIA epistatic metrics in the absence of non-motif sequence effects.** We plot the distribution of GIA epistatic metrics for discovered motifs. Class labels correspond to the interaction type for which the motif had maximum interaction effect size. For visibility purposes, we show the absolute value for each metric and clipped outliers at 0.75.

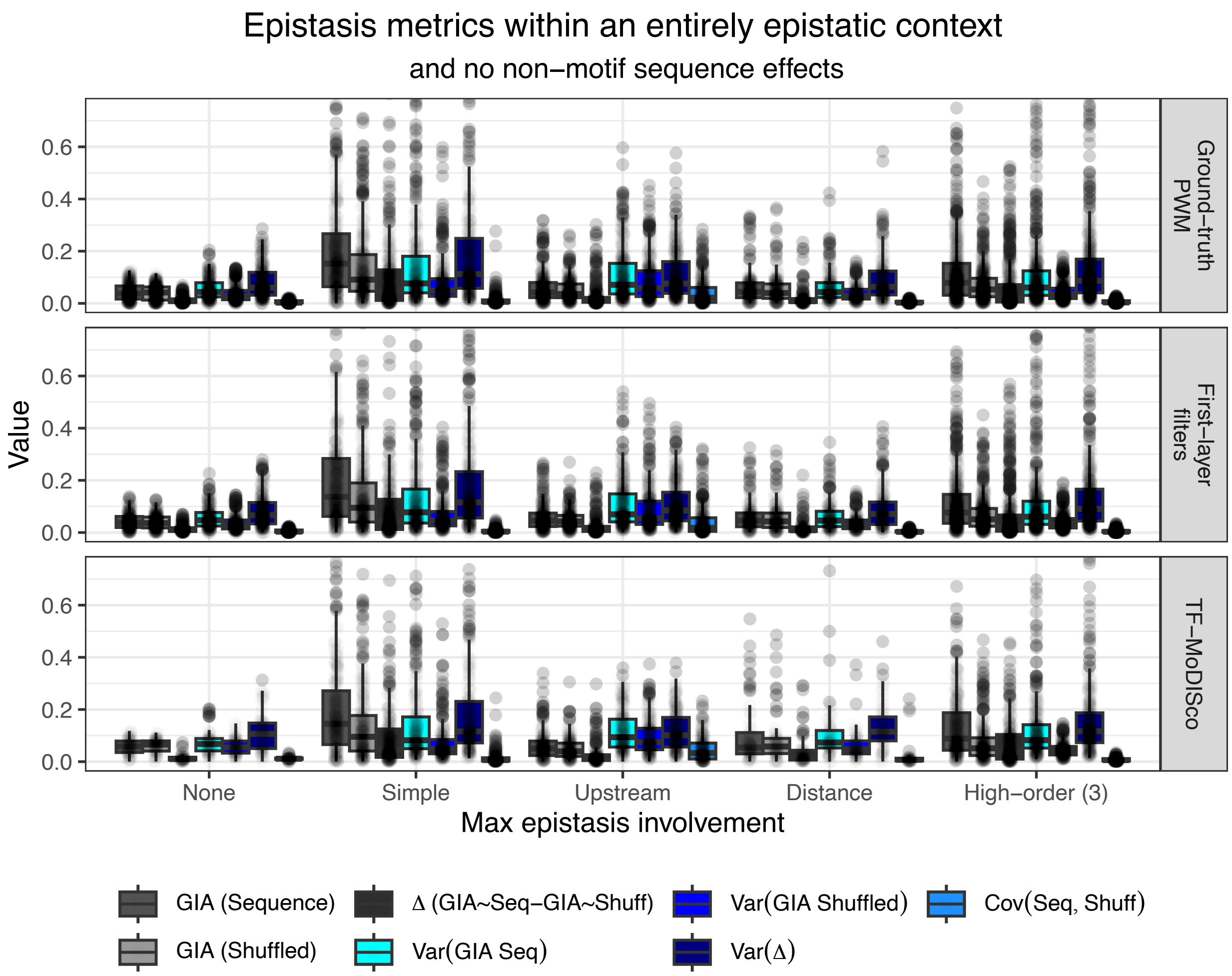


Figure S23: **GIA epistatic metrics in the presence of non-motif sequence effects.** We plot the distribution of GIA epistatic metrics for discovered motifs. Class labels correspond to the interaction type for which the motif had maximum interaction effect size. For visibility purposes, we show the absolute value for each metric and clipped outliers at 0.75.

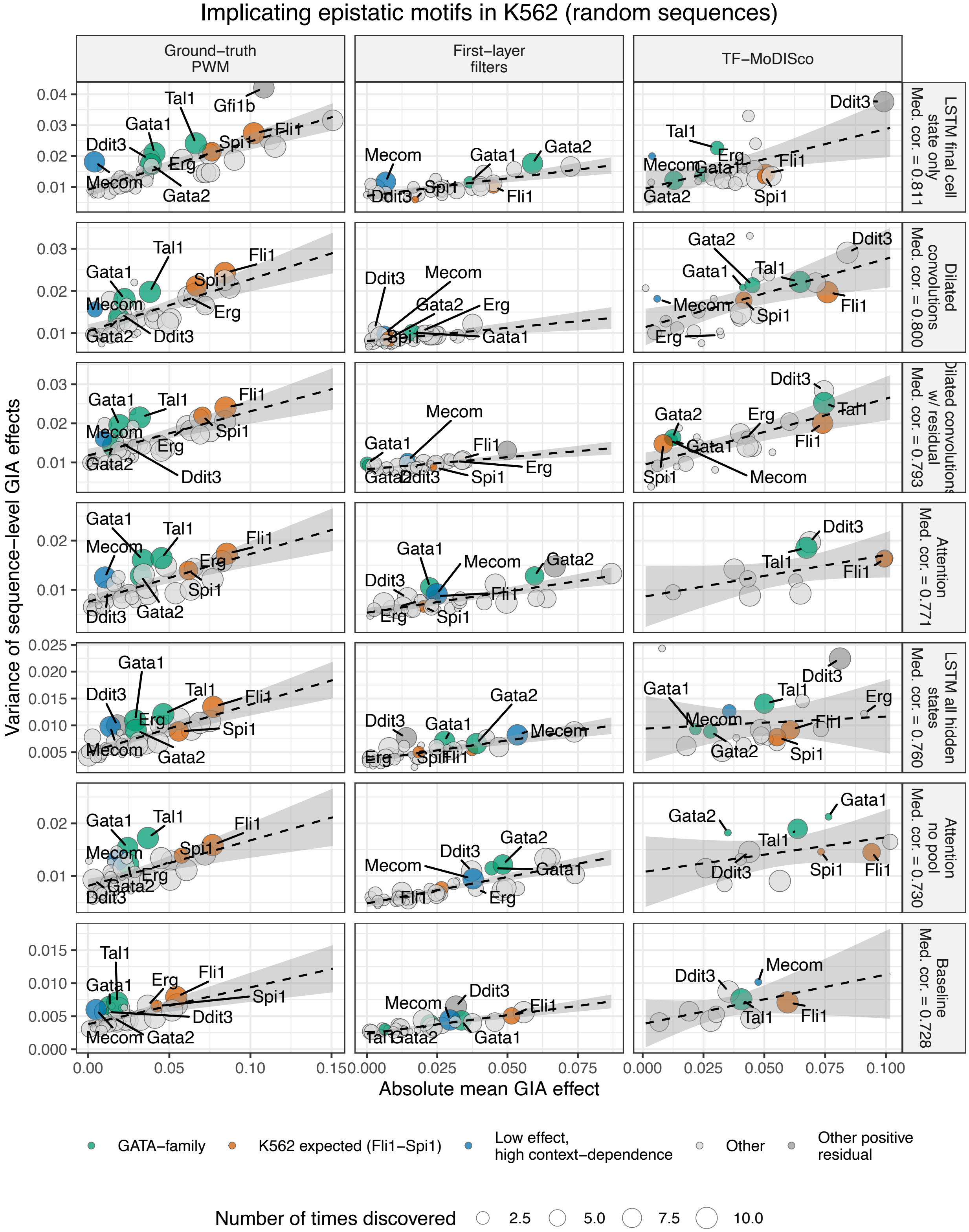


Figure S24: **Examining epistatic motifs in the K562 random sequences.** To elucidate potentially epistatic motifs, we examined the variance in the importance scores as a function of the GIA main effect.

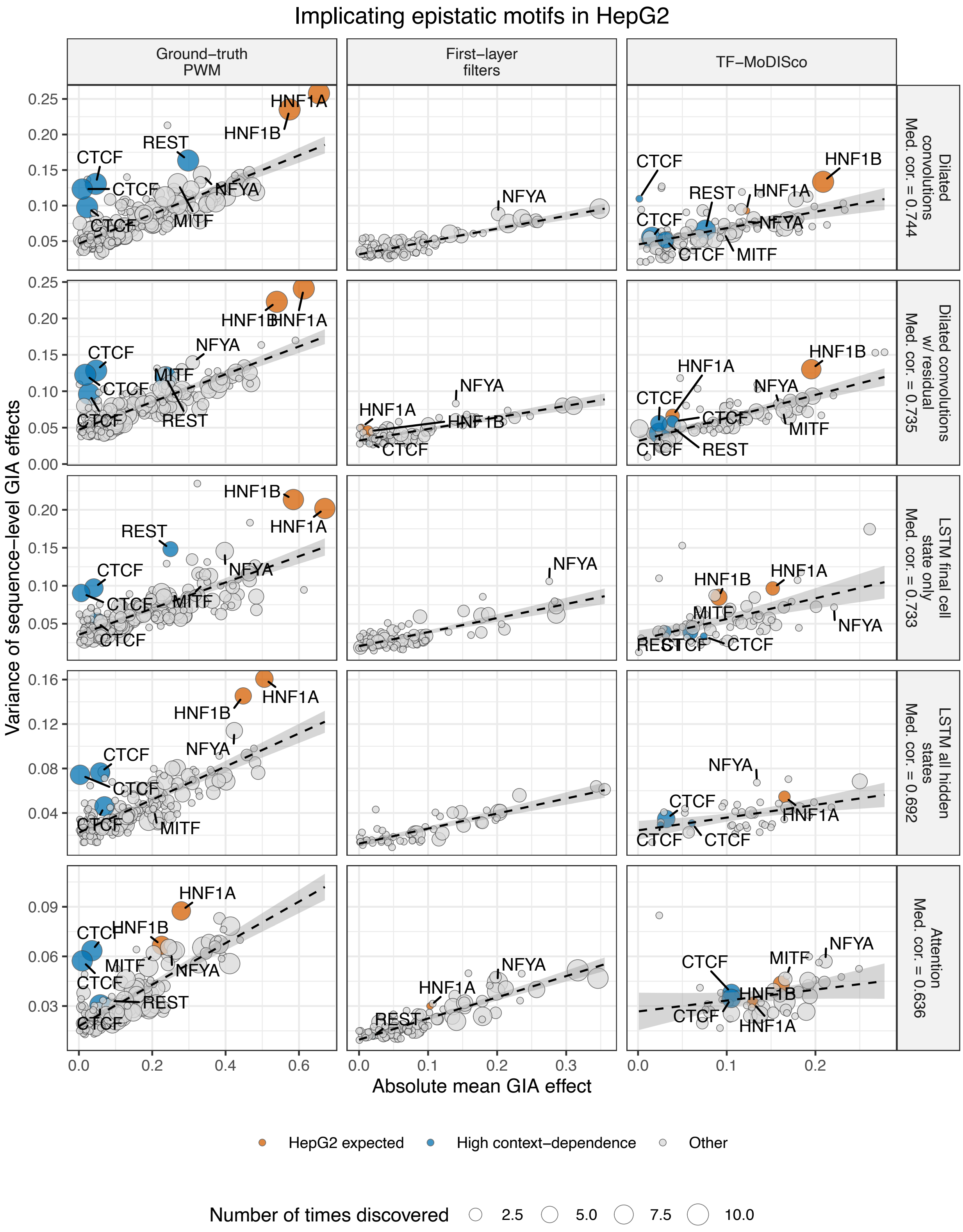


Figure S25: **Examining epistatic motifs in the HepG2 genomic-context sequences.** To elucidate potentially epistatic motifs, we examined the variance in the importance scores as a function of the GIA main effect.

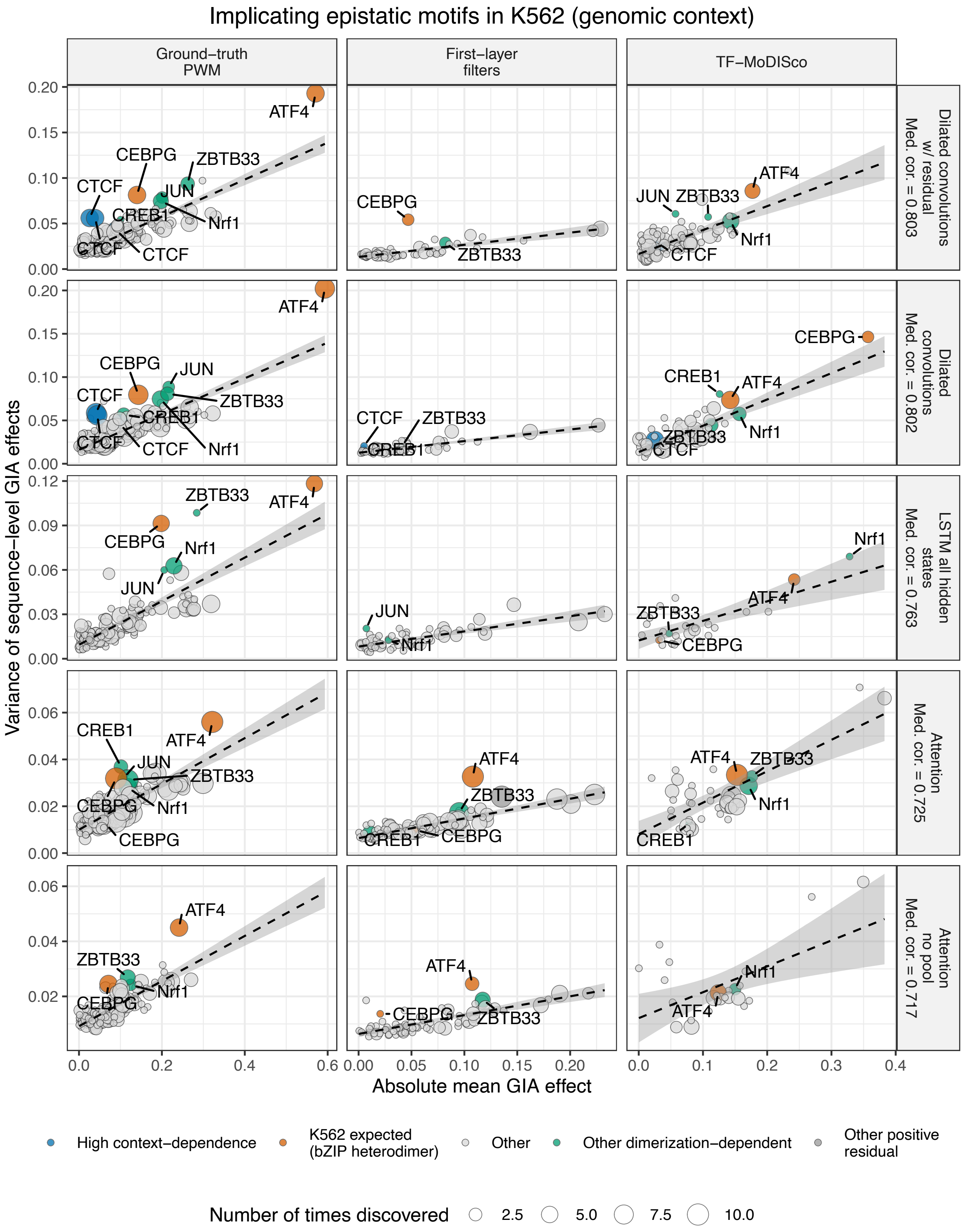


Figure S26: **Examining epistatic motifs in the K562 genomic-context sequences.** To elucidate potentially epistatic motifs, we examined the variance in the importance scores as a function of the GIA main effect.
